# A memory-based theory of emotional disorders

**DOI:** 10.1101/817486

**Authors:** Rivka T. Cohen, Michael J. Kahana

## Abstract

Learning and memory play a central role in emotional disorders, particularly in depression and posttraumatic stress disorder. We present a new, transdiagnostic theory of how memory and mood interact in emotional disorders. Drawing upon retrieved-context models of episodic memory, we propose that memories form associations with the contexts in which they are encoded, including emotional valence and arousal. Later, encountering contextual cues retrieves their associated memories, which in turn reactivate the context that was present during encoding. We first show how our retrieved-context model accounts for findings regarding the organization of emotional memories in list-learning experiments. We then show how this model predicts clinical phenomena, including persistent negative mood after chronic stressors, intrusive memories of painful events, and the efficacy of cognitive-behavioral therapies.

## Introduction

Over 70% of adults will experience traumatic stress at some point in their lifetime, including threatened or actual physical assault, sexual violence, motor vehicle accidents, and natural disasters (Benjet et al., 2016). Following trauma exposure, 20-32% of adults suffer distressing, intrusive memories of the event, hyperarousal, avoidance of event reminders, and persistent negative mood (C. R. Brewin, Andrews, Rose, & Kirk, 1999). Trauma-exposed adults also commonly experience negative mood, decreased motivation, and disrupted sleep or appetite (Goenjian et al., 2001). Historically, the Diagnostic and Statistical Manual of Mental Disorders (DSM) has divided these symptoms into categories of posttraumatic stress disorder (PTSD) or major depressive disorder (MDD), according to the prominence of memory versus mood disturbances, respectively (American Psychiatric Association, 2013). Although memory intrusions are not among the diagnostic criteria for MDD, patients with depression also report high rates of intrusive memories, replete with sensory and reliving experiences (Reynolds & Brewin, 1999). Conversely, although PTSD is not considered to be a mood disorder, affective symptoms are a diagnostic criterion and 50-75% of patients have co-occurring MDD (K. T. Brady, Killeen, Brewerton, & Lucerini, 2000). The high comorbidity and symptom overlap between these disorders has led to a search for transdiagnostic processes (Insel et al., 2010; Sanislow et al., 2010) that hold promise for the development of more efficient and efficacious treatments. Here we develop a transdiagnostic, computational model of memory in emotional disorders. This model describes how distressing events interact with the memory system, as well as the mechanisms that produce the negative mood and intrusive memories that cut across both depression and PTSD.

Research implicates learning and memory as transdiagnostic processes underlying emotional disorders (Beck, Rush, Shaw, & Emery, 1979; Bower, 1981; C. R. Brewin, 2006). Patients with PTSD experience intrusive memories of traumatic events (Ehlers, Hackmann, & Michael, 2004), but patients with depression, social anxiety, and other emotional disorders also report high rates of vivid, intrusive memories of painful events (C. Brewin, 2010; Day, Holmes, & Hackmann, 2004; Hackmann, Clark, & McManus, 2000; Muse, Mcmanus, Hackmann, Williams, & Williams, 2010; Osman, Cooper, Hackmann, & Veale, 2004; Price, Veale, & Brewin, 2012; Reynolds & Brewin, 1999; Speckens, Hackmann, Ehlers, & Cuthbert, 2007). In addition, patients with anxiety and depression tend to recall negative events more frequently than positive events, implicating mechanisms of *mood-congruent recall* (Matt, Vazquez, & Campbell, 1992; P. C. Watkins, Mathews, Williamson, & Fuller, 1992). Here, mood-congruent recall refers to better recall for memories whose emotional valence matches the current emotional context. Further, during anxiety treatment, patients benefit most when therapeutic interventions take place in an emotional context that will match subsequent retrieval contexts, suggesting processes of *emotion-state dependent recall* (M. Craske et al., 2008).

Despite these findings, current theories of learning and memory in emotional disorders do not describe whether and how mood-congruent recall, emotion-state dependent recall, and intrusive memories of painful events influence one another (C. R. Brewin, 2014; Rubin, Berntsen, & Bohni, 2008; Brown, Zandberg, & Foa, 2019). Leading theories of memory in PTSD and other emotional disorders have proposed that the arousal and valence of an experience influence its tendency to result in intrusive memories (C. R. Brewin, 2014; Rubin et al., 2008; Rubin, Dennis, & Beckham, 2011). Valence refers to the attractive (“positive”) or repellent (“negative”) properties of a stimulus, and arousal refers to the emotional intensity or degree of activation of the sympathetic nervous system due to a stimulus (Barrett, 1998; Russell, 1980; M. M. Bradley, Greenwald, Petry, & Lang, 1992; T. F. Brady, Konkle, Alvarez, & Oliva, 2008), in human memory and in emotional disorders.

However, these theories differ regarding whether high negative arousal during encoding results in intrusive memories by impairing (C. R. Brewin, 2014) or enhancing (Rubin et al., 2008, 2011) a new memory’s association to its encoding context. Further, these approaches disagree on whether it is specifically *negatively-valent* emotional arousal that results in intrusive memories, by disrupting memory-context associations during encoding (C. R. Brewin, 2014), or whether it is the intensity of emotional arousal that leads to intrusive memories, regardless of the event’s negative or positive properties, in addition to factors like the frequency of rehearsal and centrality to the person’s life story (Rubin et al., 2008, 2011). Resolving these questions is vital to improved understanding of how painful memories operate in and contribute to emotional disorders, and thereby, for generating new treatment avenues.

Computational models can help answer these questions. By building a model of the human memory system, we can test how our theory of human memory behaves under different simulated circumstances. If our model can produce the behavior we observe in the real world, then this contributes evidence in support of the theory. However, if our model cannot produce patterns that match real-world behavior, this sheds light on whether, and how, we need to change the theory. Computational modeling is especially useful when developing a theory of unobservable processes, such as what cognitive operations could generate the memory patterns that human subjects and researchers *can* observe. For example, memory researchers used computational models to develop the theory that associations between events and their contexts guide recall in human *episodic memory*, or memory for personally experienced events (Howard & Kahana, 2002a; Lohnas, Polyn, & Kahana, 2015; Healey & Kahana, 2016; S. M. Polyn, Norman, & Kahana, 2009; Talmi, Lohnas, & Daw, 2019; Moscovitch, Cabeza, Winocur, & Nadel, 2016). In the resulting *retrieved-context theory* (Howard & Kahana, 2002a; Lohnas et al., 2015; Healey & Kahana, 2014; S. M. Polyn et al., 2009; Talmi et al., 2019), the brain creates networks of associations between the contents of a memory and its context, or the internal or external surroundings (the time or place, the person’s emotional or physiological state) that were present just before or while the event took place. These contextual cues then reactivate their associated memories, and in turn, recalling a memory reactivates a mental representation of its associated contextual elements.

Here, we draw upon insights from prior clinical theories of memory in emotional disorders (C. R. Brewin, 2014; Ehlers & Clark, 2000; Foa & Kozak, 1986; Rauch & Foa, 2006) and join them with computational modeling techniques of emotion in memory (Bower, 1981; Talmi et al., 2019) to develop a new theory of the mutual influence of memory and mood in emotional disorders. We then use this model to simulate the human memory system and test whether our updated theory of emotion in episodic memory can account for the persistent negative mood and intrusive memories observed in clinical settings.

### From List Events to Life Events

We sought to develop a model of memory for life events grounded in the established scientific literature on memory for list events. Early memory scientists embraced the list-learning technique as it afforded precise experimental control over the conditions prevailing during encoding and retrieval, as well as item properties and their relations (Crowder, 1976; Kahana, 2012). Free recall of word lists constitutes one of the most widely studied list-learning paradigms — one that has generated textbook phenomena and fueled the development of major theories. In each trial, subjects study a list of sequentially presented words which they subsequently attempt to recall after a delay. This task earns the moniker “free” as the experimenter neither supplies specific retrieval cues nor imposes any requirement as to the order of recall. Because subjects must recall a specific list of items, they must direct their memory search to the items encoded in the target list-context. This context includes both the time-window in which the item occurred (temporal context) and the semantic properties shared by the items in that particular list. Each list item constitutes a mini-event, or episode, in the subject’s life, and the subject’s task is to search memory for the episodes occurring in a particular temporal context (the list). Tulving (1972) referred to recall of such episodes, bound to a specific temporal context, as *episodic memory*.

We suggest that the free recall task can serve as a building block in a model of autobiographical memory. In the real world, a complex event consists of a list of stimuli, encoded against the backdrop of other recent internal and external stimuli. We often reflect back on our experiences, either recalling them to ourselves, or to our friends. Such reminiscence is akin to free recall, and recording of these reminiscences in memory will recursively shape the way we encode and retrieve related experiences in the future. Occasionally, memories arise spontaneously, or reflecting on a given event will conjure up memories of other related events. In a free recall task, a subject will sometimes remember an item that had not occurred on a target list. Such memory intrusions, and their mnemonic sequelae, will figure prominently in our modeling of non-voluntary, unwanted memories in PTSD.

To develop our model, which we call CMR3, we used a list-learning paradigm. The list-learning paradigm is especially useful for modeling intrusions due to its ability to capture two apparently distinct processes that both share the episodic memory system: voluntary memory (strategic, in response to a targeted memory search) and involuntary memory (spontaneous and unintended). Some theorists have proposed that each type of memory should take place through distinct mechanisms; however, it is increasingly understood that both voluntary (strategic) and involuntary autobiographical memories operate through the same episodic memory system (Berntsen, 2010). Specifically, whereas people may generate additional mental context cues to guide the intended retrieval of memories, their voluntary recall is still determined by their current context. Autobiographical memories that may be experienced as “spontaneous” are also reactivated by the same process of encountering an associated spatial, temporal, emotional or other related contextual cue. Accordingly, when patients with PTSD report experiencing apparently-spontaneous memory intrusions, typically further investigation reveals the presence of one or more associated temporal, spatial, perceptual or other context cues associated to the trauma that cued the memory (Ehlers & Clark, 2000).

The externalized free recall paradigm is well suited for modeling intrusive involuntary recall because it operationalizes how both voluntary (strategic) and involuntary (spontaneous) recall outputs occur when the episodic memory system is engaged. During an externalized free recall task, subjects are instructed to report all words that enter their mind, regardless of whether or not they are trying to recall the words. In addition to the recall outputs that they report during a typical delayed free recall task, here subjects also report all other spontaneous recalls that occur during retrieval, even when subjects are aware that this spontaneous background noise is not from the intended target context for recall. Thus, intrusions are actually the result of a spontaneous, unintended process that arises as a byproduct during the subject’s intentional retrieval search. In this paradigm, intrusions occur as an unintended and undesired process that subjects must try to censor as they maintain their focus on their desired mental content (i.e., the correct results of the recall search). The two main types consist of prior-list intrusions (PLI’s), which are items the subject encountered in a prior list and does not wish to recall, and extra-list intrusions (ELI’s), which are items the subject never encountered in the current laboratory session, but has encountered previously in their life prior to the lab, and which they also do not wish to recall. In this way, ELI’s can also be conceptualized as a type of prior-list intrusion, just stemming from a prior “list” that took place prior to the laboratory session.

The list-learning paradigm is deceptively simple. Yet, it has generated a wealth of classic findings about human memory, including the primacy and recency effects (Murdock, 1962), temporal contiguity (Kahana, 1996) and semantic clustering effects (Romney, Brewer, & Batchelder, 1993; Bousfield, 1953). After their discovery in list-learning tasks, researchers have observed and replicated these classic findings in human memory for real-world events (Jansari & Parkin, 1996; Loftus & Fathi, 1985; Moreton & Ward, 2010; Uitvlugt & Healey, 2019; Healey, Long, & Kahana, 2019). As with any model, the goal of list-learning tasks is not to perfectly replicate the complexity of real-world events, but rather, to create a tractable version, or representation, of such events that draws upon the same processes that occur in memory for real-world events. This enables the experimenters to begin the process of model design.

### Retrieved-context and Episodic Memory

In retrieved-context models, the context that is encoded in association with a memory guides the later activation of that memory and determines whether it will be consciously retrieved (Healey & Kahana, 2016; Howard & Kahana, 2002a; Lohnas et al., 2015; S. M. Polyn et al., 2009; Talmi et al., 2019). Each memory’s perceptual features, such as sights and sounds, are encoded in a network of associations with the memory’s context, such as time of day, physical location, and emotions felt during the event. Later, encountering a similar context cues retrieval of the associated memories in the stored network, activating the memories whose encoded contexts have the highest overlap with the context that is present at the time of retrieval. Once a memory is retrieved, it is reencoded in association with the new context into which it was re-introduced.

Talmi et al. (2019) extended earlier retrieved-context models to account for the influence of arousal on recall performance. Their eCMR model conceptualized emotion as having a single dimension, the presence or absence of emotional arousal. In eCMR, the presence or absence of emotion is a component of memories and their contexts, and the presence of arousal strengthens the associations between items and their contexts. The resulting model, eCMR, captures key features of emotional memory in laboratory settings (Talmi et al., 2019). However, eCMR has limited ability to describe memory in emotional disorders. First, eCMR treats emotion as either present or absent, without having negative or positive valence. In addition, eCMR resets prior learning at the end of each encoding list. Thus, this model cannot distinguish between recalls from a target vs. non-target prior context, a necessary ability for modeling memory intrusions (Lohnas et al., 2015), which are of interest to accounting for intrusive trauma-memories.

### Overview

Here we propose a retrieved-context model of emotion and memory (CMR3) that considers the role of intrusions in the dynamics and persistence of affective states across a lifetime of simulated memories. First, we use a comparative modeling approach to determine the representation of emotional valence in the episodic memory system (Experiment 1, Simulation 1). Then, we demonstrate CMR3’s ability to account for mood-congruent recall (Simulation 2) and emotion-state dependent recall (Simulation 3). We then test CMR3’s ability to account for the effects of environmental negative events on negative mood (Simulation 4) and clarify the model’s predictions that repeated negative events will have a greater likelihood of becoming activated as intrusive, involuntary memories due to being associated with a wider variety of cueing contexts (Simulation 5). Then, we show the model’s ability to capture the effectiveness of positive event scheduling, a core component of behavioral activation therapy for depression (Simulations 4-5, Treatment and Post-Treatment Periods). We demonstrate CMR3’s ability to predict that high emotional arousal during encoding will lead to the development of intrusive memories (Simulation 6), and the moderating role of emotion dysregulation (Simulation 7). We show the model-predicted role of negative emotional arousal in generating heightened nowness and vividness of distressing-memory intrusions (Simulation 8), and then show CMR3’s ability to capture the efficacy of prolonged exposure therapy for alleviating intrusive memories of high-arousal negative events (*in vivo exposure*, Simulation 6; *imaginal exposure*, Simulation 9). We conclude with a discussion of retrieved-context theory’s novel predictions and its relation to current theories of memory in emotional disorders.

### A Retrieved-context Model of Memory and Mood

According to retrieved-context theory, when people form memories of new experiences, they encode the memory in association with its contextual information, such as the time and place of the experience, the thoughts and emotions present during the experience, and other internal states. Later, these contexts – such as revisiting a location where an event took place, or having an emotion that was present during the event – can cue recall of associated memories. Once context cues a memory’s retrieval, the memory reactivates its associated contexts, thus reinstating thoughts or emotions that were present during the original experience. In addition, context includes a mental representation of the perceptual features of an event, which are integrated into a composite with other contextual features. Thus, the CMR family of models supports both recall stemming from the direct perceptual features that comprised an event, such as seeing the face of someone who was present during the event, as well as other perceptual and contextual features that become cues of that memory by virtue of their shared contexts. The network of associations formed between items and past contexts can also encode semantic relationships among items, as items that share meaning tend to occur in similar temporal and semantic contexts (Howard & Kahana, 2002b; S. M. Polyn et al., 2009). Together, these episodic (contextual) and semantic (conceptual) associations guide memory retrieval.

Our model introduces two new mechanisms that improve on prior models. First, our model enables memories and their contexts to have negative, positive, and neutral emotional properties (emotional valence). Second, our model enables emotional learning to accrue over a lifetime of experiences. This also allows the model to distinguish between recall of a memory from an intended (target) context versus an unintended (non-target) context, thus allowing the model to distinguish between voluntary and intrusive (spontaneous, non-voluntary) memories. The ability to make this distinction is critical in characterizing the role of memory in the development and treatment of PTSD.

### Model Description

Here we provide a formal description of our model (Figure 1), cast in terms of equations that define the representation of items and the mechanisms that result in memory storage and retrieval. Following earlier formulations of retrieved-context theory, we assume a multidimensional feature representation of items, each denoted by **f**_*t*_ and a multidimensional representation of context that evolves as a result of each new experience and the memories it evokes (we denote the context at time t as **c**_*t*_). Our approach inherits the same equations for updating context, associating context and items, and determining recall dynamics as Polyn et al. (2009). We also inherit the additional mechanisms added by Lohnas et al. (2015) to simulate the continuity of memory across lists. We follow Talmi et al. (2019) in modeling emotion as a component of each item vector which then integrates into context. Then, we advance the model by allowing emotion to be not only present or absent as in eCMR, but further, to have positive or negative valence. In addition, we allow learning to accumulate over the course of multiple lists. This has two desirable properties: first, it allows memory to accrue over the course of long periods of time as in CMR2 (Lohnas et al., 2015), rather than a single experimental list. Second, it provides a method of operationalizing intrusive, non-voluntary memories, a symptom of great interest in PTSD and other emotional disorders. Within the list-learning paradigm described above, memory-intrusions are modeled as instances of recalling an item from an undesired, non-target list context (Zaromb et al., 2006; Lohnas et al., 2015). A model that resets the memory system at the end of each individual list cannot differentiate between target (intended) and non-target (unintended, undesired) recall contexts, and thus cannot model recall of a memory-intrusion from a non-target recall context. We call this updated model CMR3 and present a mathematical description below.

**Figure 1.**
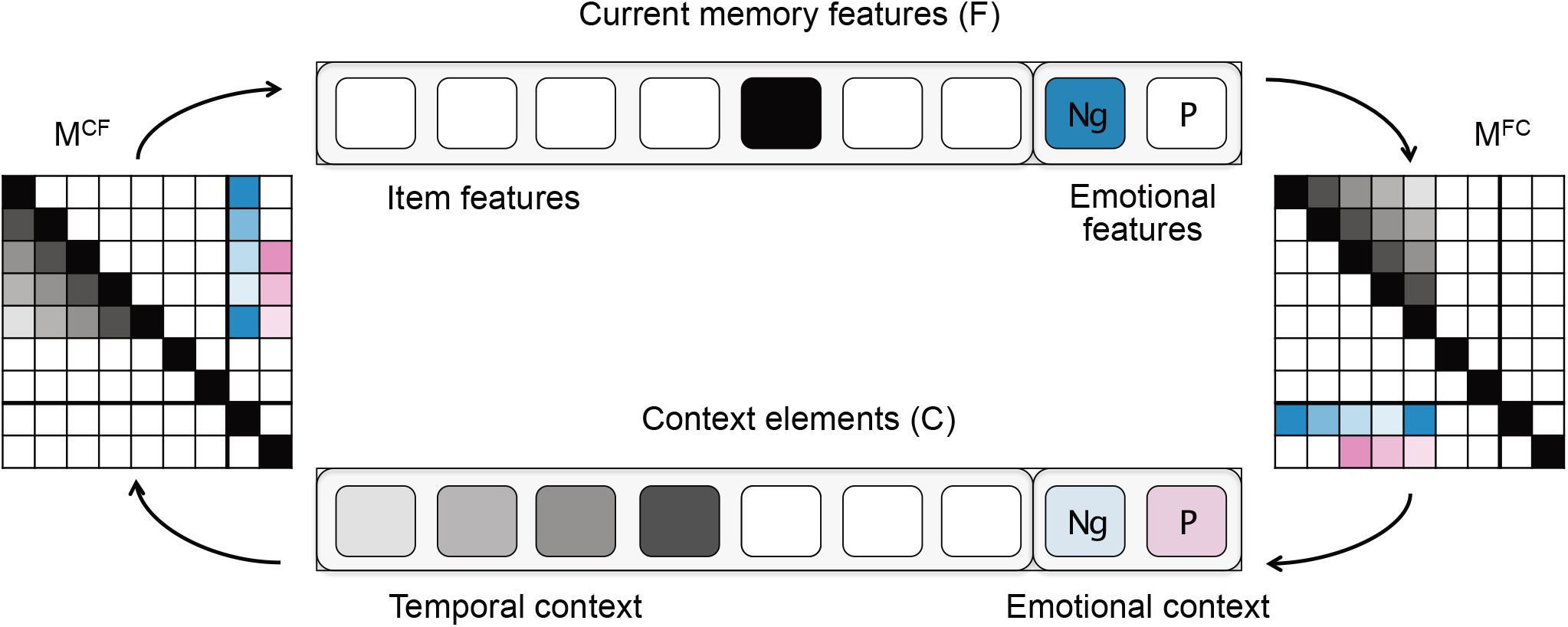
A Retrieved Context Theory of Memory and Emotion. The vector F represents event features comprising both item and emotional content. Item features are either present (have a value of1.0, shown in black) or absent (have a value of 0.0, shown in white). Valence is encoded by setting either the Ng cell (where values are shown in blue) or the P cell (where values are shown in pink) to 1.0, indicating negative and positive valence, respectively. Values of 0.0 for both cells, shown in white, indicate neutral valence. The vector C represents context and comprises recent event features(temporal context) and previously evoked emotions (emotional context). The shaded colors represent the decay of context features, which can take any value between 0.0 and 1.0. The F and C vectors interact through two weight matrices, *M*^*CF*^ and *M*^*FC*^, which store the strengths of associations between event features and context elements. These associations are also shown in shaded colors to indicate values between 0.0 and 1.0. Context cues memories for recall by activating context-to-item associations stored in *M*^*CF*^. New events update context with their own features, and recalling a memory reactivates its associated context, via item-to-context associations stored in *M*^*FC*^. See text for details.

#### Item Representation

In CMR3, each element of **f**_*t*_ represents whether that item is present or absent in the current memory episode, and each element of **c**_*t*_ represents the extent to which prior memories or cognitive states are still active during encoding. In addition to the representations of item features in **f**_*t*_ and temporal context in **c**_*t*_, Polyn et al. (2009) introduce an additional subregion in each vector to contain source-memory attributes, such as the encoding task conducted during learning (S. Polyn, Norman, & Kahana, 2009) or the presence of emotion (Talmi et al., 2019). In eCMR, emotion is represented as simply present or absent, taking a binary value of 0 or 1 in the source-memory subregion. To model how memory differentially contributes to negative or positive mood, we updated the emotional subregion of both feature and context vectors to contain two cells. In **f**_*t*_, one cell indicates whether the item is negative, the second whether it is positive, and neutral items have no content in either cell. In **c**_*t*_, one cell holds the accumulation of negative context and the other holds the accumulation of positive context. We chose two separate cells due to findings that negative and positive emotion may operate via separate cognitive systems (Cacioppo & Berntson, 1994; P. J. Lang, 1995).

#### List Representation

As in CMR2 (Lohnas et al., 2015), we concatenated all presented lists in both the feature and context vectors, to allow the model to carry information forward that was learned in prior lists. Each list is separated by the presentation of disruptor items. These disruptor items induce a shift in temporal context between lists, but they do not form associations with other items in the Hebbian weight matrices, as is the case during the presentation of true items. The Hebbian weight matrices store the associations between item features and context features that form during the experiment (see *Encoding* section, below). We begin each simulation by presenting an initial disruptor item so that the initial context vector is not empty (Lohnas et al., 2015). In addition, we present CMR3 with a disruptor item after each encoding list to model the distractor task used in Experiment 1.

#### Context-Updating

During encoding, as each item is presented, it enters the current context, causing other contextual elements to decay. Thus, context evolves according to the equation:

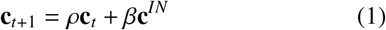

As such, context becomes a recency-weighted sum of past context states. The features from **f**_*t*_ that will enter into the new context, represented by the term **c**^*IN*^, are determined by multiplying the weight matrix of item-to-context associations, which we call *M*^*FC*^, by **f**_*t*_, the vector of current features, and then norming the resulting vector, such that 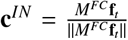. In equation 1, *ρ* scales the magnitude of prior context so that **c** does not grow without bounds (see Howard & Kahana, 2002a). The *β* parameter determines the rate at which new temporal context enters the updated context vector. This rate differs depending on whether the subject is encoding a new event (*β*_*enc*_), retrieving a new memory (*β*_*rec*_), registering a shift between list contexts (*β*_*post*_), or experiencing a context shift due to a distractor task (*β*_*distract*_). Emotional context is stored in the source region of the context vector, and it updates according to the same equation that governs updating for temporal context, but at the rate of *β*_*emot*_. The parameter *β*_*emot*_ is the parameter termed *β*_*source*_ in Talmi et al. (2019), renamed here to emphasize that it governs the updating of emotional source context.

#### Emotional Context and the Calculation of “Mood”

As described above, CMR3 represents emotional context in the source region of the context vector. As in item features (see subsection “Item representation,” above), one cell accumulates negative emotional context, and one cell accumulates positive emotional context. The model updates both types of emotional valence at the same rate of *β*_*emot*_, but the values in each cell can vary independently of one another, depending on both the emotional valence of new events and on the emotional context reactivated by recalling a memory. Thus, the model produces independent indices and separate activations of positive and negative emotion. However, to simplify graphical representation of the model’s predicted emotional responding in our simulations, we created a composite measure of “mood.” We calculate “mood” as the difference between the degree of positive context and negative context that is present on a given timestep (*mood*_*t*_ = *c*_*pos,t*_ − *c*_*neg,t*_. We used a simple difference metric due to keep values on an intuitive scale between -1 (completely negative emotion) and 1 (completely positive emotion), with 0 representing equivalent levels of positive and negative emotional responding.

#### Encoding

As each newly presented item evolves and updates cognitive context, its features form associations with the elements of context present during encoding. The memory and context representations, **f**_*t*_ and **c**_*t*_, interact via two Hebbian associative (outer-product) weight matrices, which model the strength of associations from the studied items to their encoding context, *M*^*FC*^, and from context to associated items, *M*^*CF*^. Because **f**_*t*_ and **c**_*t*_ each have two subregions – the first devoted to individual features of the memory, or item features, and its temporal context, and the second consisting of two cells devoted to emotional valence (see item representation), *M*^*FC*^ and *M*^*CF*^ have corresponding subregions. In the upper-left quadrant, each matrix contains associations between item features, **f**_*items*_, and temporal (or non-emotional) context elements, **c**_*temp*_. In the lower-left quadrant of *M*^*FC*^ and the upper-right quadrant of *M*^*CF*^, each matrix contains the associations between item features and emotional context elements, **c**_*emot*_, as below:

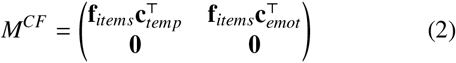

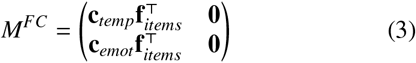

Prior to the experiment, we initialize each weight matrix as an identity matrix of rank *i* + 2, where *i* is the total number of items presented in the experiment. Two is the number of elements contained in the emotional subregion of the feature and context vectors. In the process, the subregions containing associations between items’ temporal features and emotional context are initialized to zero. The pre-experimental Hebbian matrices, called 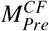 and 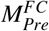, are then scaled by (1 − *γ*_*CF*_) and *M*^*CF*^ by (1 − *γ*_*FC*_), respectively. The parameters *γ*_*CF*_ and *γ*_*FC*_ are the learning rates for context-to-item and item-to-context associations, respectively (see Learning Rates, below).

#### Semantic Associations

To model semantic effects on recall, we construct a matrix of inter-item semantic associations, called *M*^*S*^. CMR3 represents longstanding semantic associations between pairs of items by adding *M*^*S*^ to the upper-left quadrant in 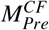, which contains associations between items’ temporal features and temporal context. Each entry in the *M*^*S*^ matrix equals the dot product similarity between vector representations of each item’s features (S. M. Polyn et al., 2009). We define these similarities using Google’s Word2Vec algorithm (Word2Vec; Mikolov, Chen, Corrado, & Dean, 2013), which uses the co-occurrence of different words across multiple texts to determine vector representations of each word. The similarity between a pair of items can then be calculated using the dot product between each pair of item vectors. When adding *M*^*S*^ to the initialized *M*^*CF*^, we scale *M*^*S*^ by the parameter s. Thus, s determines the degree to which semantic associations guide the retrieval process.

#### Learning Rates

New episodic associations accumulate as different memory features and contextual elements are active at the same time. We preserved the learning rule from CMR (S. M. Polyn et al., 2009), in which memories form associations with the context at the current time step, rather than at the preceding timestep. Thus, as each new memory is encoded, the change in associations stored in each weight matrix equals:

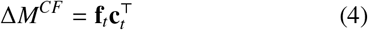

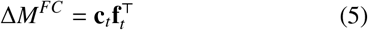

The rate at which these associations integrate into the *M*^*FC*^ and *M*^*CF*^ weight matrices is determined by three parameters: *γ*_*FC*_, *γ*_*CF*_, and *γ*_*emot*_, which is the parameter termed 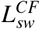 in Talmi et al. 2019. The parameters *γ*_*CF*_ and *γ*_*FC*_ are the learning rates of new context-to-item and item-to-context associations, respectively. When updating associations in *M*^*CF*^, *γ*_*emot*_ is the learning rate of new associations between item features and emotional context in *M*^*CF*^, which allows memories to form new associations with emotional vs. non-emotional context at different rates. The learning rate in the other quadrants is set to 0.0, since these associations do not contribute to the encoding and retrieval processes in the current model equations (S. M. Polyn et al., 2009; Talmi et al., 2019).

The *M*^*FC*^ matrix handles the degree to which recalled memories reactivate their associated temporal and emotional (source) contexts. For simulating mood in emotional disorders, we were primarily interested in the overall reactivation of emotional context during recall, as an operationalization of mood, rather than in the relative activation levels of temporal and emotional context elements. Therefore, for simplicity and to conserve free parameters, we allowed associations between item features and both types of context in *M*^*FC*^ to form at the same rate, *γ*_*FC*_. However, in future work, a more complex version of the model could allow these associations (between item features and temporal context versus emotional context) to accumulate at separate rates in *M*^*FC*^. Because source features in F do not currently contribute to the recall process, the upper- and lower-right quadrants of *M*^*FC*^ are set to 0.0 (S. M. Polyn et al., 2009).

Thus, before Δ*M*^*FC*^ and Δ*M*^*CF*^ integrate into *M*^*FC*^ and *M*^*CF*^, we scale each Δ*M* elementwise by its respective matrix of learning rates, *L*^*FC*^ and *L*^*CF*^:

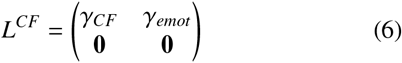

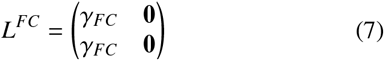

Following Sederberg et al (2008), CMR3 models increased attention to early list items by scaling each value in equation (4) by 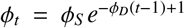, where *ϕ*_*S*_ scales the overall level of the primacy effect and *ϕ*_*D*_ determines the rate of decay in this primacy effect, as the *i*th item is presented. Thus, at a given point in the experiment, the strength of the associations between items and contexts stored in each weight matrix are given by the equations:

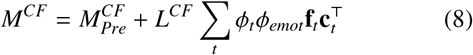

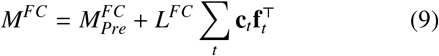

As in Talmi et al. (2019), we include a parameter, *ϕ*_*emot*_, which modulates the strength of context-to-item associations when an emotional item evokes arousal during encoding. For items that evoke no arousal, *ϕ*_*emot*_ takes a value of 1.0, and for items that evoke emotional arousal, *ϕ*_*emot*_ takes a value greater than 1.0 (Talmi et al., 2019).

#### Recall

The state of context at the moment of retrieval serves to activate its associated memory elements. This takes place by multiplying *M*^*CF*^, by the current context vector at retrieval, **c**_*R*_. In the resulting vector of item activations, each item is activated to the extent that its context during encoding is similar to the context that is present at the start of the retrieval process, such that the vector of item activations is given by:

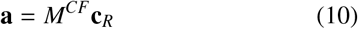

Thus, each memory’s features are activated according to the strength of the dot-product similarity between the context vector present during its encoding and the context that is present during retrieval, as well as the strength of preexisting semantic associations. The activated elements then enter an evidence accumulation process, in which the evidence for the retrieval of any given item is driven by its activation value in **a**. On each step of this leaky accumulator process (Usher & McClelland, 2001), the vector of evidence, **x**, evolves according to the following equation:

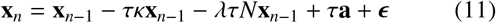

Accordingly, the evidence on the current time step equals the level of evidence on the last step (**x**_*n*−1_) minus the decay in that evidence over time, minus lateral inhibition from other activated items (N is a matrix with 1’s along the diagonal and -1’s at all other entries, such that the summed activation of other items is subtracted from the given item’s activation, and then scaled by *λ* and the time-decay scalar, *τ*), plus incoming item activations, and plus noise, *E*. When the evidence in **x**_*n*_ for an item *i* passes a certain threshold, *θ*_*i*_, that item emerges as a candidate for recall. For computational efficiency, we limit this evidence accumulation race to the items with the highest activations (*N*_*items*_ = 2× list length), since the other items are very unlikely to emerge as successful candidates for recall (Lohnas et al., 2015).

The winning item then updates the **f**_*t*_ vector with its features, meaning that the **f**_*t*_ vector once again holds a 1 in the cell corresponding to the retrieved item, and 0’s in the other items’ cells. The item’s associated context is then retrieved by multiplying *M*^*FC*^ by the newly updated **f**_*t*_ vector. If the recalled item’s evoked context, **c**^*IN*^, is sufficiently similar to the current context (**c**^*IN*^ · **c**_*t*_ *>* **c**_*thresh*_), with **c**_*thresh*_ being a threshold scalar), then that item is reinstated into **f**_*t*_. Since the CMR3 model context vectors represent two subregions, each of which is updated and normed separately, the total contextual similarity value ranges between 0.0 and 2.0. Whether or not the retrieved item is reinstated, it cues its associated context, which evolves the context vector according to the same equations that update **c**_*t*_ during encoding. Once an item has been retrieved, the threshold of evidence needed to subsequently recall this just-retrieved item is equal to the threshold *θ*_*i*_ = 1 + *ωα*^*j*^, where *j* represents the number of times that a different item has been subsequently retrieved. Here, *α* = [0.0, 1.0), such that *ω* raises the absolute level of the threshold, and *α* decays this newly raised threshold each time the given item is reactivated. All dynamics of this part of the recall process (the reinstatement of items after the leaky accumulator) follow the model equations defined in CMR2 (Lohnas et al., 2015).

#### Correspondence between CMR3 and Prior Retrieved-context Models

The model we formalize above, CMR3, builds on earlier retrieved-context models of episodic memory (Howard & Kahana, 2002a; S. M. Polyn et al., 2009; Lohnas et al., 2015; Kahana, 2020). The CMR model extended the original temporal context model (Howard & Kahana, 2002a; Sederberg, Howard, & Kahana, 2008) by including semantic and source information into the latent representation of context (S. M. Polyn et al., 2009). Talmi et al. (2019) added (negative) arousal as an additional contextual attribute, and additionally allowed arousal to modulate the learning rate for context-to-feature associations. The CMR2 model (Lohnas et al., 2015; Healey & Kahana, 2016) extended the basic model to account for learning and interference effects across multiple lists, as well as providing a post-retrieval recognition mechanism that plays an important role in monitoring potentially intrusive memories. CMR3 extends CMR2 by adding a multivalent representation of emotion, allowing for neutral, positive, negative, or mixed emotional states. CMR3 extends eCMR by incorporating the multilist capabilites of CMR2.

### Experiment 1: The Role of Emotional Valence in Memory

To understand how episodic memory can promote negative versus positive emotion in emotional disorders, it is crucial to understand how emotional valence is represented in and evoked by human memory. When freely recalling lists of studied items, subjects tend to recall clusters of negative, positive, and neutral items (Long, Danoff, & Kahana, 2015). This *emotional clustering effect* suggests that not only the presence of emotion, but also its valence, guides the organization and retrieval of items in memory. Here, we replicate the emotional clustering effect in an independent dataset (Kahana, Aggarwal, & Phan, 2018; Aka, Phan, & Kahana, 2020) (described as Experiment 1, below).

Subsequently, we evaluate the ability of three variants of retrieved context theory to model this phenomenon: one in which there is no representation of emotional information other than as part of the memory’s semantic content (Lohnas et al., 2015), one in which a single binary feature indicates whether a memory does or does not possess emotional information (Talmi et al., 2019), and a third in which two binary attributes separately indicate whether a memory possesses positive or negative emotional attributes (the current CMR3 model). We fit each of these models to the emotional clustering effect along with a set of other benchmark recall phenomena, and to the emotional in free recall (see Simulation 1).

## Method

Ninety-seven young adults (Mean age = 22; 51.5% female) completed delayed free recall of 576 unique lists, each comprising 24 unique words drawn from a subset of 1638 nouns from the University of South Florida word pool (D. L. Nelson, McEvoy, & Schreiber, 2004). Following list presentation and before the start of the recall period, subjects performed a 24 second arithmetic distractor task intended to attenuate recency sensitive retrieval processes. During the 75s recall period, subjects attempted to vocally recall as many words as they could remember, in whatever order they came to mind. Because subjects must recall words from the most recent list, this task measures the ability to localize memory to a specific context (i.e., this specific list, in this specific session). Because subjects may recall the study words in any order (i.e., freely), the order in which these words are recalled suggests how the items were organized in memory. For full details of the experiment stimuli and methods see Kahana et al. (2018).

We assessed the valence of each word using ratings from a prior norming study (Long et al., 2015). In this rating study, 120 subjects on Amazon’s Mechanical Turk (MTURK; Mason & Suri, 2012) generated valence ratings for each item, rating them on a scale from 1-9, where 1 indicated negative valence and 9 indicated positive valence. Words were defined as ’negative’ if their valence rating fell below 4, ’neutral’ if between 4-6, and ’positive’ if above 6. The resulting valences of the 864 words in the experiment word pool were 25.6% positive, 8.9% negative, and 65.5% neutral.

## Results

Data from this experiment served both to replicate the emotional clustering effect reported by Long et al. (2015) and to obtain individual subject parameter sets for use in Simulations 2-9. By having each subject contribute 24 sessions of trials (totalling 576 lists) the present study provided sufficient data to allow for stable modeling at the individual subject level. To evaluate the effect of emotional valence on recall organization we conducted a conditional probability analysis of transitions among items according to their positive, negative, or neutral valence. Following Long et al. (2015), we adjusted probabilities to account for different rates of negative, positive, and neutral items in each study list. We then examined subjects’ tendencies to transition between same-valent or differently-valent items, with higher probabilities for same-valent items indicating a higher tendency to cluster these items together during recall. We also assessed five benchmark recall phenomena unrelated to the emotional valence of items: serial position and output order effects, contiguity effects, semantic clustering effects, and patterns of prior- and extra-list intrusions (Kahana, 2020). We explain each of these effects below.

The serial position curve (Figure 2A) illustrates the probability of recalling items as a function of their ordinal position within the study list. As is typical of delayed recall experiments, this analysis revealed a strong primacy effect – superior recall of early list items – but little or no evidence of recency – i.e., enhanced recall of final list items. The primacy effect seen in the serial position curve largely reflects subjects increased tendency to initiate recall with early list items, as seen in the probability of first recall (PFR) curve shown in Figure 2B. The lag-conditional response probability (Lag-CRP) shows the probability of successively recalling items as a function of their separation (lag) in the study list. This curve illustrates the contiguity effect, wherein subjects exhibit a strong tendency to successively recall items studied in neighboring list positions (small values of interitem lag) and they make these transitions with a forward bias (see Figure 2C.

**Figure 2.**
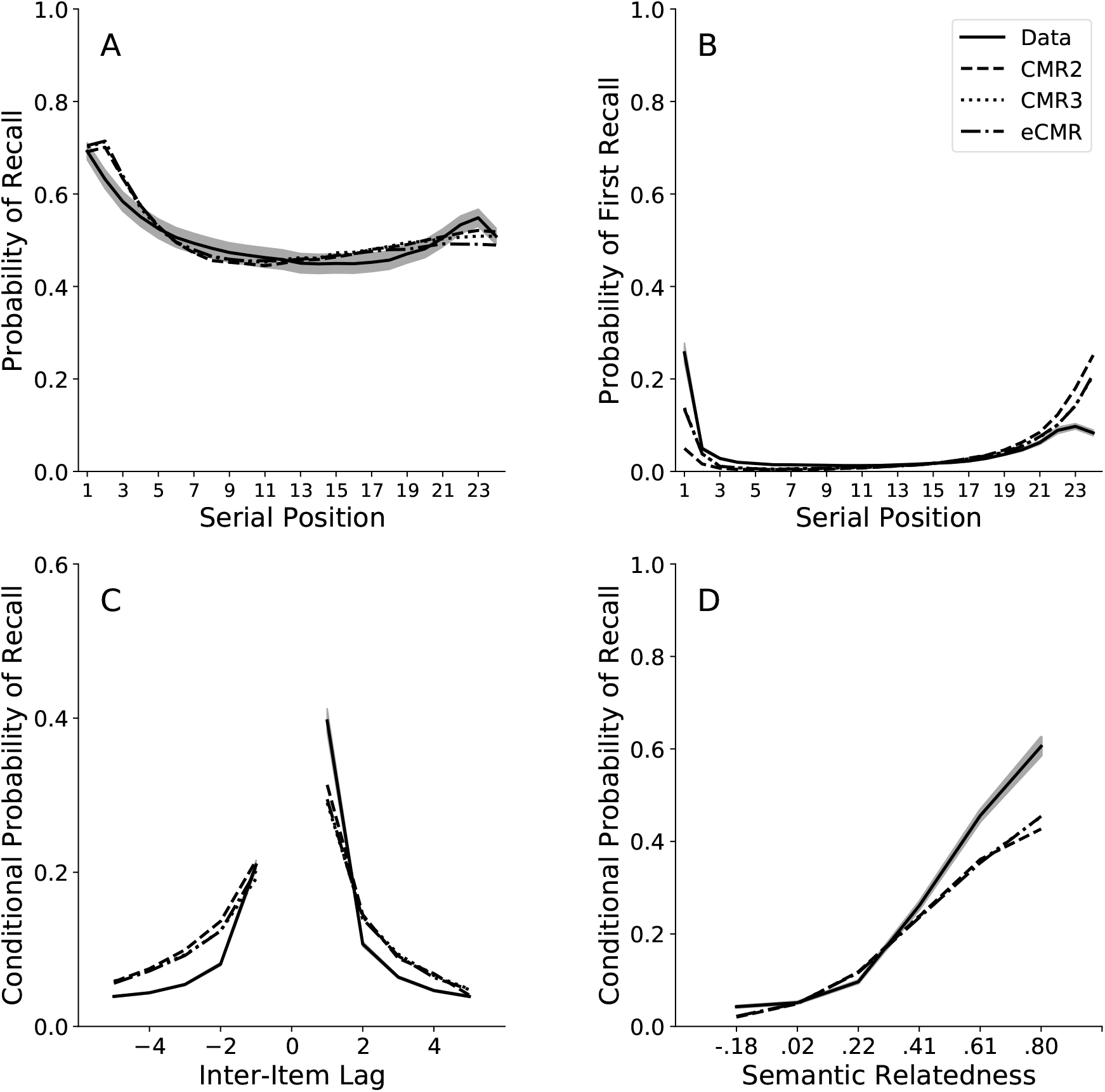
Modeling the Organization of Memories during Free Recall. We fit CMR2, eCMR, and CMR3 to serial position effects and to temporal, semantic, and emotional clustering in a delayed free recall task. Error regions represent ± 1 *S*.*E*.*M*., calculated across subjects (data). Error regions across parameter sets’ simulated dataset values are omitted for visibility, but see Table 3 for RMSE fit indices. **A**. Recall probability as a function of items’ presentation order during encoding. **B**. The probability that an item will be recalled first, as a function of an items’ presentation order during encoding. **C**. The probability that items will cluster together during recall according to their proximity to one another during encoding. For an example, the value at a lag of +2 is the probability that, given the recall of item A, the item that is recalled directly after item A was encoded 2 positions after item A during study. **D**. The probability that items will cluster during recall according to their semantic similarity to one another. The curve depicts the probability that, given the recall of item A, the item recalled right after item A has a certain level of similarity with A, plotted for six semantic word2Vec bins (-.18 to .02, .02 to .22, .22 to .41, .41 to .61, .61 to .80, .80 to 1.0).

We used a similar conditional probability analysis to measure subjects’ tendency to successively recall semantically related list items (*semantic clustering*; Howard & Kahana, 2002b; Long et al., 2015). To represent semantic relations among the words, we used Word2Vec, which is a method of estimating the representations of words in vector space, or word “embeddings” (Mikolov et al., 2013). Word2Vec identifies the semantic meaning of a word based on what other words accompany it across many different settings in which it is used. Using machine-learning, Word2Vec trains on a database of online texts and then generates vectors representing the dimensions of each word based on the other words that typically accompany it. The inter-item similarity for each pair of words is then calculated as the dot-product between each word-vector in the pair. Using Word2Vec, we generated inter-item similarities for each pair of words in the 1,638 noun word pool, from which we computed six bins of similarity values (-.18 to .02, .02 to .22, .22 to .41, .41 to .61, .61 to .80, .80 to 1.0), following procedures in Howard and Kahana (2002b) and Long et al. (2015). Figure 2D shows that subjects exhibited a semantic clustering effect, successively recalling semantically related items with far greater probabilities than unrelated items.

In addition, we calculated the average rate at which subjects mistakenly recalled items from lists preceding the target list (*prior-list intrusions*, or PLI’s), and mistaken recalls of items that were never presented during the experiment (*extra-list intrusions*, or ELI’s). As expected based on previous research (Kahana, Dolan, Sauder, & Wingfield, 2005; Zaromb et al., 2006), our healthy young adult subjects produced low rates of PLI’s and ELI’s per session, with means of 0.10 (SEM = .01) PLI’s and 0.34 (SEM = .03) ELI’s across subjects.

As in Long et al. (2015), we observed a small, but significant tendency to recall items in clusters of the same emotional valence (Figure 3). That is, upon recalling a negative item, subjects were more likely to next recall another negative item than a positive item, and upon recalling a positive item, subjects were more likely to next recall another positive item than a negative item (Figures 3). To test the significance of this effect, we conducted a three (Initial Item Valence: Negative, Positive, or Neutral) × three (Transition-Item Valence: Negative, Positive, or Neutral) linear mixed effects model predicting transition probabilities (lme4 R package; Bates, Mächler, Bolker, & Walker, 2015). We set neutral valence as the base category, and we used a random intercept to account for the dependency of observations within each subject. The fixed-effects estimates and significance testing using the Satterthwaite’s method for approximating degrees of freedom (lmerTest R package; Kuznetsova, Brockhoff, & Christensen, 2017) are presented in Table 1.

**Figure 3.**
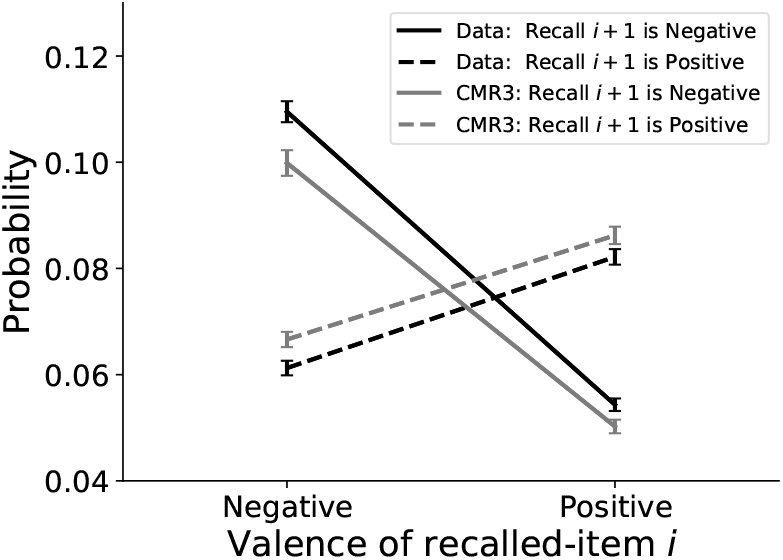
The Emotional Clustering Effect. In a free recall task, subjects tend to successively recall items that share contextual features, including their emotional valence. Thus, after recalling a negative (positive) item they tend to recall other negative (positive) items. CMR3 provides a good fit to this effect, as seen in the conditional probability of transitions among items as a function of their valence. Solid lines represent the tendency to transition to a negative item, and dashed lines represent the tendency to transition to a positive item.

**Table 1.**
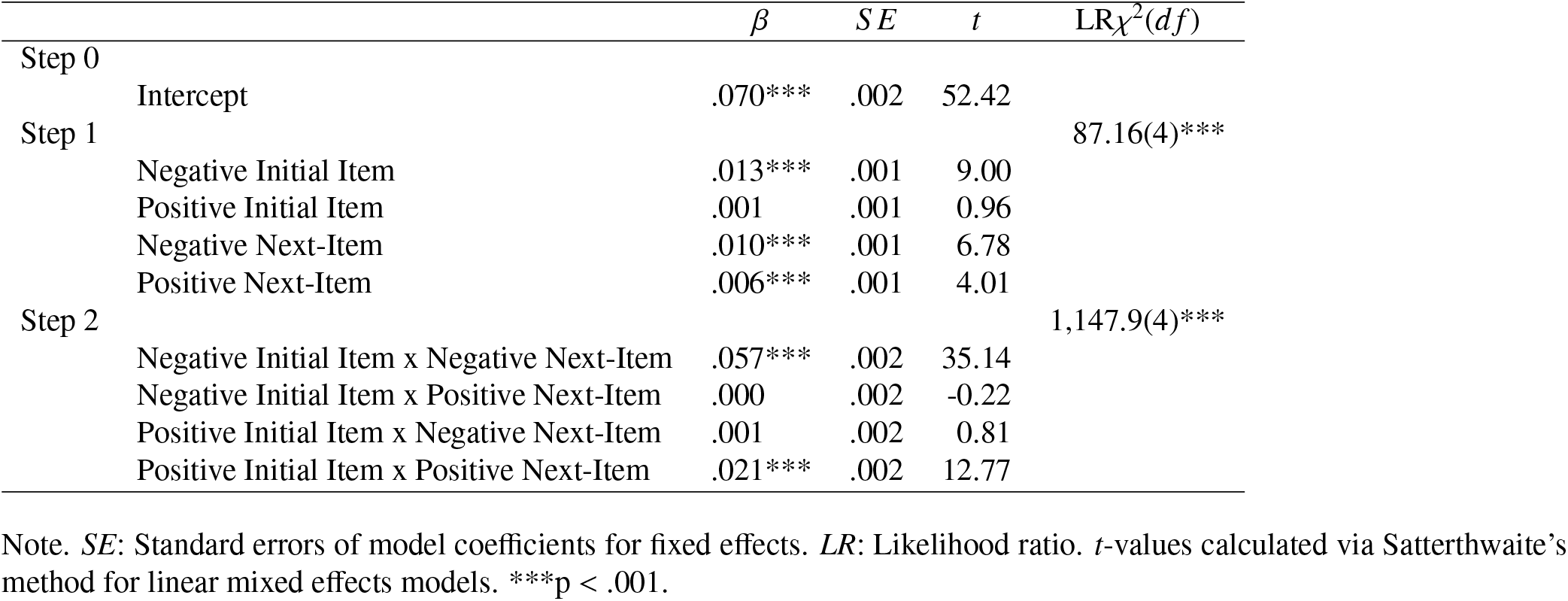
Transition Probabilities Regressed on Initial-item Valence and Next-item Valence.

The resulting interactions indicated that negative items were significantly more likely to be recalled together with other negative items than with non-negative items, *β* = .057, *t*(768) = 35.1, *p <* 0.001, and positive items were significantly more likely to be recalled together with other positive items than with non-positive items, *β* = .021, *t*(768) = 12.8, *p <* 0.001. Taken together, our results replicate findings that positive and negative items tend to cluster together with same-valent items during recall (Long et al., 2015). Next, we use a computational approach to identify which representation of emotion in memory is optimal to predict this effect.

### Modeling Emotional Organization of Memories

To model the effect of emotional valence on the organization of memories, as seen through the dynamics of the free recall process, we fit CMR2, eCMR, and CMR3 to the behavioral effects observed in Experiment 1. This allowed us to compare three possible representations of emotion in memory encoding and retrieval processes. In CMR2 (Lohnas et al., 2015), we implemented emotion solely as part of items’ semantic content: that is, as a component of inter-item semantic associations. In eCMR (Talmi et al., 2019), we additionally included an item’s status as emotional versus neutral (i.e., the presence or absence of emotional content) as a part of item and context features. Because we were interested in capturing patterns of item intrusions across lists, and the original eCMR does not have multilist capabilities, we fitted eCMR with multilist capabilities as in CMR2 and CMR3. In CMR3 (see *Model Overview*), we additionally included emotional valence as part of item and context features.

### Model Specification

To isolate the effects of emotional valence, we held the arousal parameter, *ϕ*_*emot*_, constant at 1.0 for each model. At this value, arousal neither enhances or weakens the context-to-item associations in *M*^*CF*^. This allowed us to test each model’s ability to capture emotional patterns in the behavioral data solely due to how it represents emotion’s dimensions in the feature and context vectors. In addition, the words presented in the free recall tasks have emotional valence properties but likely induce minimal or absent levels of arousal during the free recall task (Long et al., 2015). As in the behavioral analyses in Experiment 1, we used Word2Vec (Mikolov et al., 2013) to generate inter-item similarities for each pair of items, which were added to the pre-experimental *M*^*CF*^ matrix that represents associations between items and contexts.

### Simulation 1

#### Method

Using a particle swarm optimization method, we fit each of the three CMR variants to data from each individual subject. To obtain the best-fitting parameters for each model, fit to each subject, we minimized the *χ*^2^ error between each model’s predictions and the actual data for that subject (see Appendix). This produced a set of 97 best-fitting parameters sets and predicted data points for each model (Table 2).

**Table 2.** Best-fitting Parameters for Each Model, Averaged across Individual-subject Fits.

|  | CMR2 |  | eCMR |  | CMR3 |  |
| --- | --- | --- | --- | --- | --- | --- |
|  | Mean | SEM | Mean | SEM | Mean | SEM |
| $\beta_{enc}$ | .298 | .008 | .295 | .008 | .286 | .007 |
| $\beta_{rec}$ | .713 | .013 | .717 | .013 | .701 | .012 |
| $\beta_{distract}$ | .458 | .018 | .481 | .020 | .453 | .020 |
| $\beta_{post}$ | .672 | .021 | .675 | .025 | .733 | .020 |
| $\beta_{emot}$ | — | — | .369 | .018 | .322 | .016 |
| $\gamma_{FC}$ | .799 | .010 | .800 | .010 | .813 | .009 |
| $\gamma_{CF}$ | .872 | .009 | .845 | .010 | .869 | .009 |
| $\gamma_{emot}$ | — | — | .394 | .025 | .417 | .026 |
| $\phi_S$ | 1.413 | .074 | 1.009 | .064 | .957 | .072 |
| $\phi_D$ | .611 | .042 | .747 | .037 | .728 | .037 |
| $\kappa$ | .257 | .012 | .279 | .013 | .293 | .012 |
| $\eta$ | .184 | .010 | .181 | .010 | .189 | .010 |
| $\lambda$ | .118 | .010 | .115 | .011 | .125 | .010 |
| $s$ | 1.044 | .041 | 1.049 | .039 | 1.117 | .049 |
| $\omega$ | 22.821 | .383 | 22.702 | .406 | 22.605 | .382 |
| $\alpha$ | .934 | .006 | .958 | .0006 | .954 | .006 |
| $c_{thresh}$ | .339 | .018 | .991 | .038 | .992 | .038 |
| $\phi_{emot}$ | 1.0 | | 1.0 | | 1.0 | |

Note. $\beta_{enc}$ = drift rate for temporal context during encoding. $\beta_{rec}$ = drift rate for temporal context during recall. $\beta_{distract}$ = drift rate for temporal context during a distractor item or shift between mental tasks. $\beta_{post}$ = drift rate for temporal context at the end of a recall session. $\beta_{emot}$ = drift rate for emotional context. $\gamma_{FC}$ = learning rate for item-to-temporal context associations. $\gamma_{CF}$ = learning rate for temporal context-to-item associations. $\gamma_{emot}$ = learning rate for emotional context-to-item associations. $\phi_S$ scales the primacy effect and $\phi_D$ determines its rate of decay over the presentation of new items. $\kappa$ = rate of evidence decay in the leaky accumulator due to the passage of time. $\eta$ = width of the normal distribution of noise values for each item's recall evidence in the leaky accumulator. $\lambda$ = degree of lateral inhibition between items in the leaky accumulator. $s$ = scales the strength of inter-item semantic associations. $\omega$ scales the raised threshold for item repetition, and $\alpha$ scales the decay in that threshold as subsequent items are recalled. $c_{thresh}$ = threshold of similarity between an item's encoding and retrieval contexts needed in order to consider the item a correct recall. $\phi_{emot}$ = a scalar that magnifies the strength of context-to-item associations during the encoding of a high-arousal negative item.

Fitting individual subjects served two purposes: First, for each model, we used the resulting sets of parameters to generate three simulated datasets, comprised of the responses predicted by each model (CMR2, eCMR, or CMR3) for each subject. The resulting parameters enable simulating an entire 24-session experiment, using the actual list items that a given subject experienced to determine the sequence of items presented to the model. The model then generates a set of simulated recall sequences, which we analyze to estimate the same behavioral patterns shown in Figure 2. For each model’s simulated dataset of “virtual subjects,” we repeated the full behavioral analyses that we ran on the empirical data. This enabled us to evaluate and compare each model’s ability to fit the aggregate behavioral data. Second, obtaining the best-fitting parameters to each subject allowed us to later examine the predictions of retrieved-context theory for how different people, whose minds operate according to different cognitive parameters, respond to negative life events with quick recovery, vs. negative mood or intrusive, non-voluntary memories (Simulations 4-9).

### Simulation Results

To assess each model’s ability to match the aggregate data, we calculated three measures of fit: (1) the *χ*^2^ goodness of fit index that was minimized while fitting the models; (2) the Bayesian Information Criterion (BIC) to account for the different number of parameters across models (Kahana, Zhou, Geller, & Sekuler, 2007; S. M. Polyn et al., 2009; Schwarz, 1978); and (3) the RMSE, to identify which specific behavioral analyses determined each model’s overall ability to fit the data (see Appendix). The resulting *χ*^2^ error values were *χ*^2^(60) = 74.1, *p* = .10 for CMR2, *χ*^2^(58) = 55.2, *p* = .58 for eCMR, and *χ*^2^(58) = 56.7, *p* = .52 for CMR3, indicating that all three model fits had non-significant error. The resulting BIC’s were -347.06 for CMR2, -345.65 for eCMR, and -353.82 for CMR3, where lower (i.e., more-negative) values represent improved model fits. The results indicate that CMR3 provided the best balance of model parsimony and error minimization, followed by CMR2 and then eCMR.

Next, we examined which behavioral effects distinguished each model’s ability to fit the aggregate data. We calculated RMSE values for each behavioral analysis (Table 3). The CMR3 model provided the smallest total RMSE, followed by CMR2, and then eCMR, where smaller values indicate better model fit. Comparing eCMR and CMR3, eCMR provided lower RMSE’s for the positive lags of the Lag-CRP and the frequency of extra-list intrusions. Conversely, CMR3 provided the lowest RMSE for the emotional clustering effect, followed by CMR2 and then eCMR. CMR2 provided worse fits to the semantic clustering effect and the probability of first recall, suggesting that the model may have had to sacrifice fits to these data in its attempts to capture emotional clustering patterns.

**Table 3.** RMSE’s for Behavioral Analyses.

|  | CMR2 | eCMR | CMR3 |
| --- | --- | --- | --- |
| Total | .064 | .061 | .058 |
| Emotional Clustering | .007 | .008 | .006 |
| SPC | .025 | .028 | .027 |
| PFR | .059 | .038 | .037 |
| Left CRP | .036 | .030 | .031 |
| Right CRP | .043 | .050 | .053 |
| Semantic CRP | .084 | .076 | .076 |
| ELI's | .325 | .322 | .325 |
| PLI's | .098 | .098 | .098 |

Note. Total: All data points. Emotional Clustering: Emotional clustering effect. SPC: Serial position curve. PFR: Probability of first recall. Left CRP: Negative lag values of the Lag-Conditional Response Probability graph. Right CRP: Positive lag values of the Lag-Conditional Response Probability graph. Semantic CRP: Semantic conditional response probability graph. ELI's: Frequency of extra-list intrusions per list. PLI's: Frequency of prior-list intrusions per list.

Visual inspection of Figure 2 confirms that, with some variation, all three models predicted the shape of the serial position curve and the pattern of recall initiation, which indicates that subjects tend to initiate recall with items from the beginning or the end of the list (Figures 2a,b). All three models also captured subjects tendency to successively recall items studied in neighboring list positions (the temporal contiguity effect) (Figure 2c) and their tendency to successively recall semantically similar items (the semantic clustering effect, (Figure 2d).

The models differed, however, in their ability to capture the emotional clustering effect, the key phenomena of interest in this simulation (Figure 3). Figure 4b displays the tendencies of each model to over-predict or under-predict item clustering according to the emotional valences of just-recalled and next-recalled items. By not including emotion as a type of episodic features and contexts, CMR2 under-predicted the tendency of same-valent items to cluster together. However, by not differentiating between items of different valences, eCMR over-predicted the tendency of opposite-valent items to cluster together. In CMR3, allowing the emotional content of items and contexts to have both negative and positive emotional valence best captured the tendency of same-valent items to cluster together and the tendency of oppositely-valent items to not cluster together.

**Figure 4.**
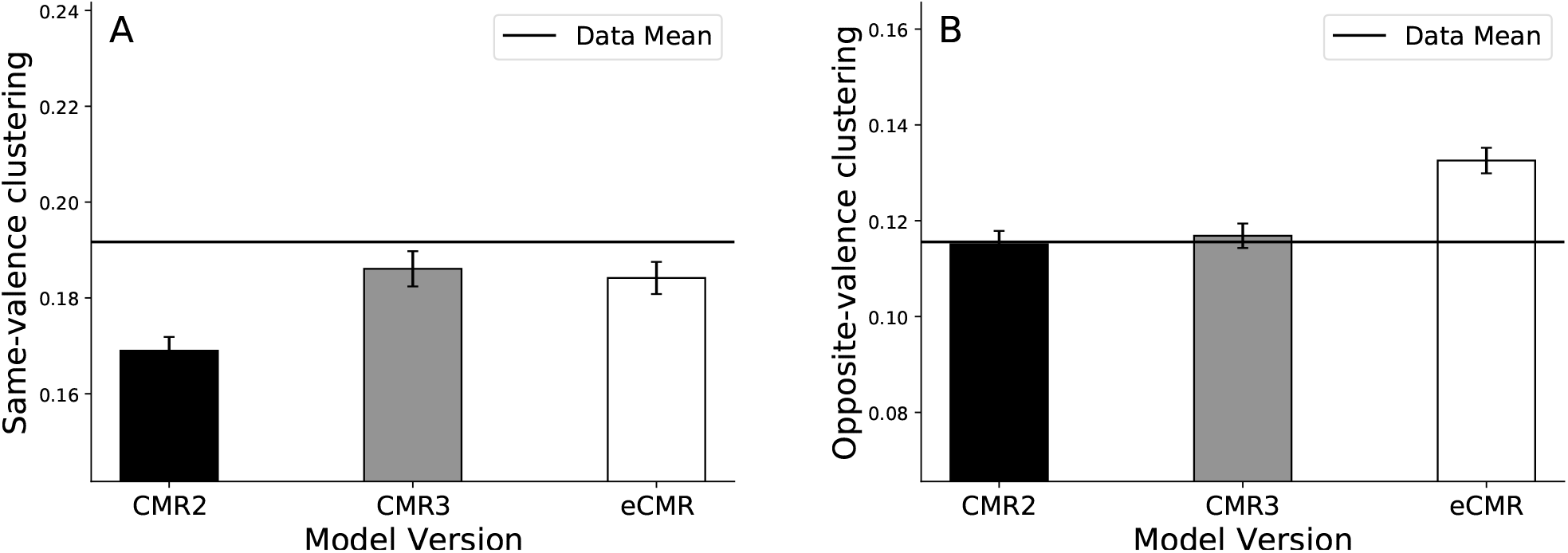
Modeling the Organization of Memories during Free Recall, Continued. The emotional clustering effect represents the tendency of items to cluster during recall according to their emotional valence. This figure shows the model-predicted degree of clustering among **A**. same-valent and **B**. oppositely-valent items for CMR2 (black bars), CMR3 (gray bars), and eCMR (white bars). Error bars represent ± 1 *S*.*E*.*M*..

We have shown that to capture the emotional clustering effect, a model should represent emotion as having positive and negative valence (as in CMR3). One of the strengths of this dataset is its ability to show how emotional valence influences recall in the absence of significant arousal. A limitation is that we did not orthogonally manipulate arousal and valence. Further, as we limited our stimuli to low-arousal items in order to isolate the effects of valence, Experiment 1 did not produce an emotionally-enhanced recall effect, which eCMR was designed to capture (Talmi et al., 2019). Testing CMR3 and the updated eCMR (expanded here to have multilist capabilities) on experiments with high-arousal items can reveal the relative influences of emotional valence and arousal in episodic memory, and would indicate whether separate modeling assumptions are required to capture enhanced recall of high-arousal items (Talmi et al., 2019). In subsequent sections, we show how CMR3 can also capture patterns of negative vs. positive mood and memory in emotional disorders.

### Mood-congruent Recall

Having shown that CMR3 can account for the emotional clustering effect without sacrificing its fit to benchmark recall phenomena, we next examined CMR3’s ability to account for mood-congruent and emotion-state dependent recall. We consider these two classic phenomena of memory and emotion as a prelude to our main objective, which is the development of a new theory that extends the retrieved context framework to account for persistent negative mood and the production of intrusive, non-voluntary memories. We see these as potential transdiagnostic processes underlying major depression and PTSD.

### Motivation

A patient in a depressive episode may readily recall memories of loss or failure, while memories of happy events elude him (Teasdale & Russell, 1983). Yet, once recovered, this patient may tend to recall positive memories and have difficulty remembering the events that occurred while they were depressed. Heightened recall for memories whose emotional attributes match the person’s current emotional state, or *mood-congruent recall*, has been observed widely across both clinical and healthy populations: e.g., for sadness and loss-related stimuli in patients with depression (B. P. Bradley, Mogg, & Williams, 1995; Teasdale & Fogarty, 1979; P. C. Watkins et al., 1992), for threat-related stimuli in patients with anxiety (Mathews, Mogg, May, & Eysenck, 1989, but see Bradley et al., 1995) and patients with PTSD (Paunovic, Lundh, & Öst, 2002), and for positive stimuli in healthy subjects (P. C. Watkins et al., 1992). Talmi et al. (2019) have previously demonstrated the ability of retrieved-context models (eCMR) to capture mood-congruent recall for emotional versus non-emotional items. Here, we test whether retrieved-context theory can account for mood-congruent recall within distinct types of emotional states (decreased recall of positive events when in a negative mood state, and vice-versa).

### Simulation 2

#### Method

We tested CMR3’s ability to predict mood-congruent recall. Specifically, we simulated the results from a series of free recall experiments in which mood-congruent recall would be expected to occur. First, we simulated 97 virtual subjects by specifying the model with each set of parameters generated while fitting CMR3 to the behavioral data in Experiment 1. Each set of parameters defines the virtual subject’s memory system, determining the responses that the model (i.e., the virtual subject) will give while completing the simulated task. Then, we presented each virtual subject with the following task. The simulated task consisted of 60 encoding lists, with each containing a randomly permuted mixture of positive (30%), negative (30%), and neutral (40%) items.

To ensure that negative, positive, and neutral items would all have equivalent semantic structures, we drew the interitem similarity for each pair of items at random from a normal distribution (*µ* = .2, *σ* = 0.5). A value of -1.0 represents maximum dissimilarity, and 1.0 represents maximum similarity. To simulate a set of unique items, no item pairs were assigned an inter-item similarity of 1.0. Finally, we set the s parameter equal to 0.15, which scaled down the inter-item semantic similarities by that value. This allowed the model outputs to more strongly reflect the episodic dynamics of interest.

In this task, each virtual subject began studying each list in a neutral state, with no emotional content in the subject’s context vector. After each encoding list, we simulated a mood induction prior to free recall, by setting the virtual subject’s emotional context to contain negative (20 lists), neutral (20 lists), or positive (20 lists) emotional content. The model then predicted the subject’s recall responses during free recall, in which emotional context was allowed to evolve naturally in response to the emotional content of the retrieved items. We then ran this simulation for each virtual subject: that is, each of the 97 sets of parameters obtained in the fits to behavioral data in Simulation 1. Across simulations, we analyzed the probability of recall for positive, negative, and neutral stimuli, conditional upon the type of mood induction prior to recall.

#### Results

Using the sets of parameters obtained in Simulation 1 (see Table 2), we simulated a set of 97 virtual subjects, by specifying the CMR3 model with each set of parameters and obtaining each model-predicted output for a virtual recall task. For an overview of which parameters we manipulate in each simulation, see Table 4.

**Table 4.**
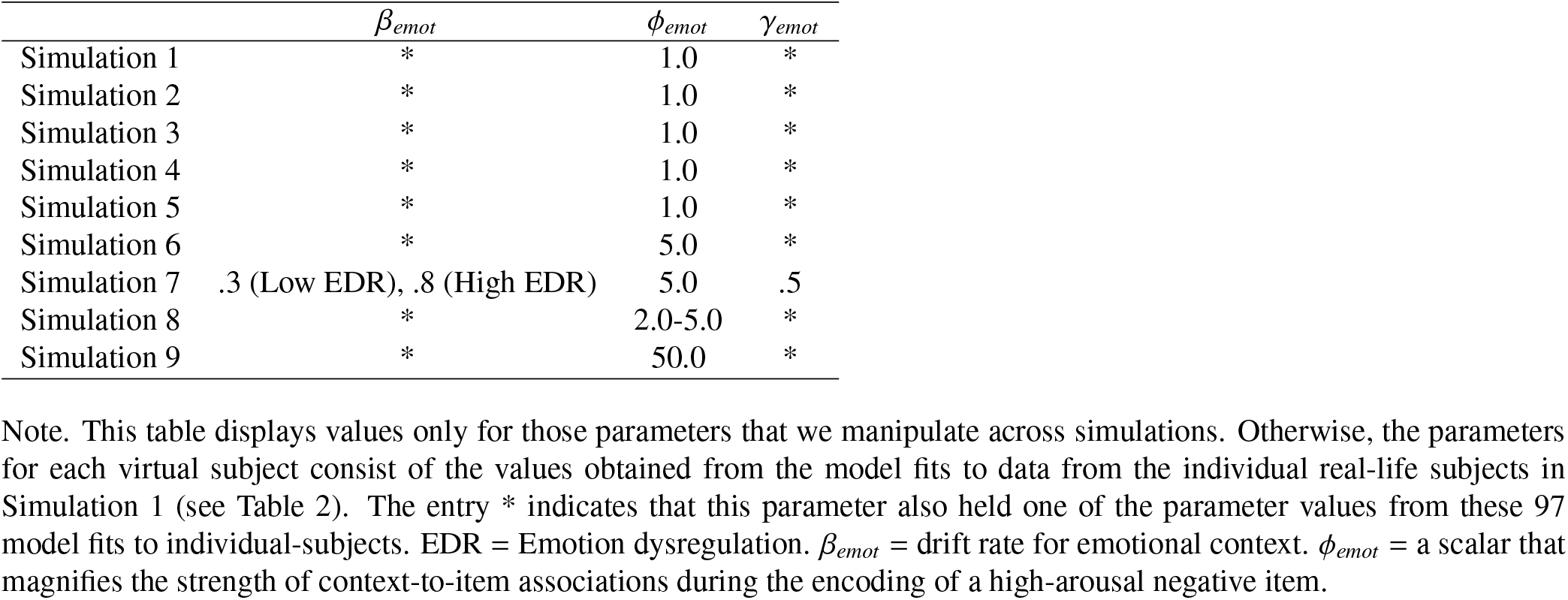
Parameter Differences across Simulations

| | $\beta_{emot}$ | $\phi_{emot}$ | $\gamma_{emot}$ |
| --- | --- | --- | --- |
| Simulation 1 | * | 1.0 | * |
| Simulation 2 | * | 1.0 | * |
| Simulation 3 | * | 1.0 | * |
| Simulation 4 | * | 1.0 | * |
| Simulation 5 | * | 1.0 | * |
| Simulation 6 | * | 5.0 | * |
| Simulation 7 | .3 (Low EDR), .8 (High EDR) | 5.0 | .5 |
| Simulation 8 | * | 2.0-5.0 | * |
| Simulation 9 | * | 50.0 | * |

Note. This table displays values only for those parameters that we manipulate across simulations. Otherwise, the parameters for each virtual subject consist of the values obtained from the model fits to data from the individual real-life subjects in Simulation 1 (see Table 2). The entry \* indicates that this parameter also held one of the parameter values from these 97 model fits to individual-subjects. EDR = Emotion dysregulation. $\beta_{emot}$ = drift rate for emotional context. $\phi_{emot}$ = a scalar that magnifies the strength of context-to-item associations during the encoding of a high-arousal negative item.

Then, we assessed the model-predicted effects of mood state during retrieval on the recall of emotionally valent and neutral items. Across parameter sets, model predicted that negative or positive mood states cue more-frequent recall of items that match the current emotional state (Figure 5a). Further, the model predicted that, during a neutral mood, emotional items were more likely to be recalled overall, with probabilities of 0.66 (SEM = .021) for negative items, 0.65 (SEM = .023) for positive items, and 0.61 (SEM = .024) for neutral items. This matches findings that in laboratory tasks lacking mood induction manipulations, and in which most subjects likely have neutral mood during the task, emotional items have a recall advantage over neutral items (Long et al., 2015; Siddiqui & Unsworth, 2011; Talmi et al., 2019). In CMR3, this occurs because studying or recalling an item evokes emotional context consistent with its emotional valence, and these emotional features serve as the input to the context updating equation. As a result, items that induce a particular emotional context during their encoding have a heightened match when the emotional context evoked by the pre-recall mood induction has matching emotional elements.

**Figure 5.**
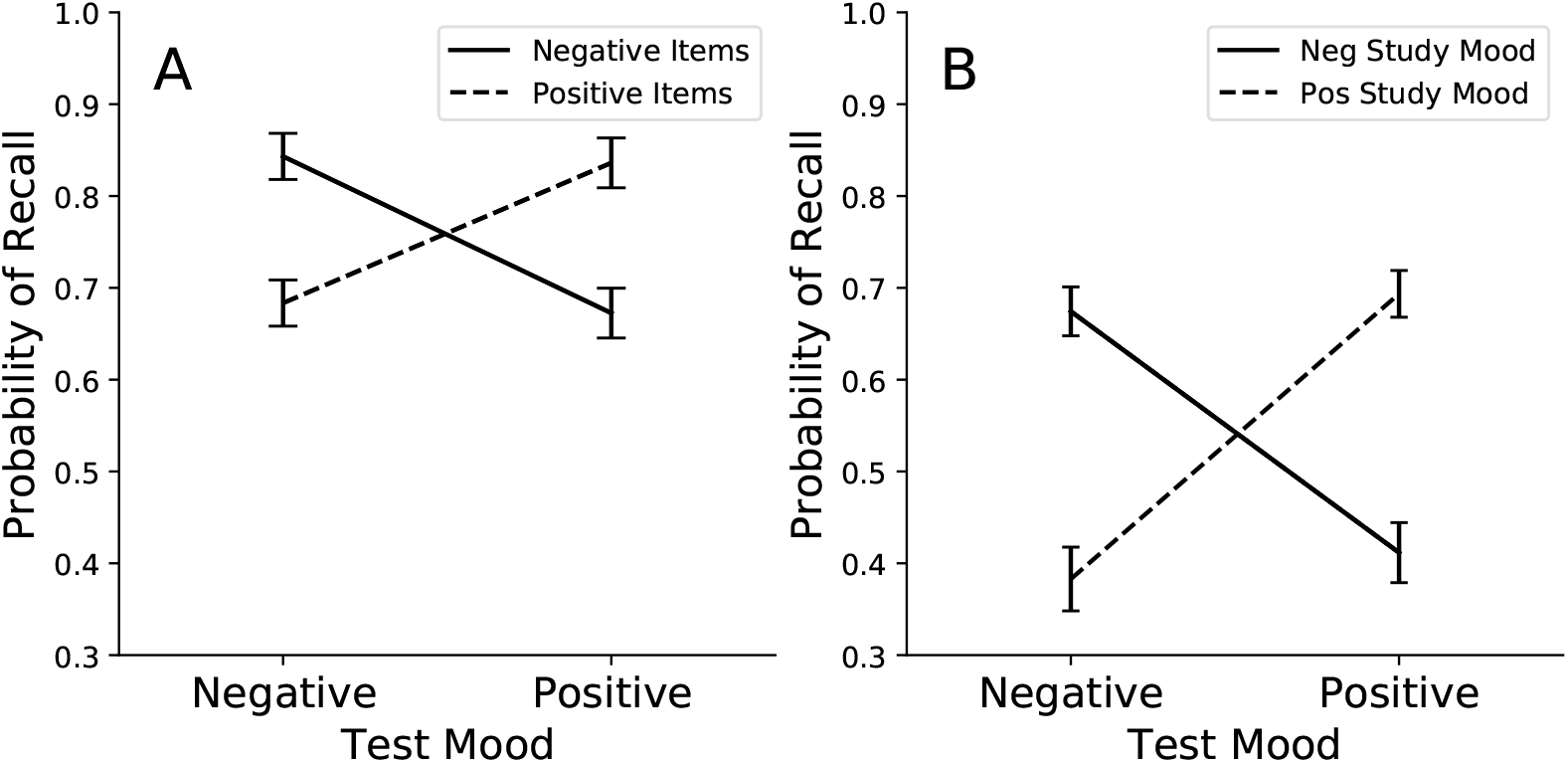
Simulating Mood-congruent and Emotion-state Dependent Recall. Using parameters obtained from the behavioral fits in Simulation 1, we used CMR3 to predict recall outputs based on two factors: (1) the match between an item’s valence and the emotional state during retrieval (mood-congruent recall), and (2) the match between the emotional state during encoding and retrieval, regardless of the item’s valence (emotion-state dependent recall). **A**. Mean probability of recalling a negative item (solid lines)or a positive item (dashed lines), depending on the valence of the recall context, which is marked on the horizontal axis. **B**. Mean probability of recalling a neutral item depending on whether it was encoded and retrieved in matching versus non-matching emotional states. The valence of emotion during retrieval, or Test mood, is marked on the horizontal axis. Solid lines represent negative mood during study, and dashed lines represent positive mood during study. Error bars represent ± 1 SEM calculated across all parameter sets’ predictions.

In CMR3, we operationalized neutral mood as the reduced or relative absence of emotional context, rather than as a distinct mood state. This produces mood-congruent recall for positive and negative mood states, and enhanced recall of emotional material in a neutral mood state, consistent with empirical findings (Long et al., 2015; Siddiqui & Unsworth, 2011; Talmi et al., 2019). Enhanced recall of emotional material in a neutral mood state emerges due to emotional context adding additional contextual elements that can cue the retrieval of these memories. However, an alternate way to operationalize neutral mood would be to consider it a “mood state” in and of itself, rather than the absence of negative or positive emotional content. In this design, the model would produce enhanced recall for neutral items in neutral mood states.

### Emotion-state Dependent Recall

#### Motivation

Whereas patients learn new skills and insights in a calm, therapeutic setting, they must often recall these skills for use in settings fraught with intense emotion. Emotion-state dependent recall refers to better recall of learned items or associations when they are retrieved in a context similar to that in which they were encoded. Emotion-state-dependent recall is a variant of the more general state-dependent recall effect, in which information learned in one environment or state is better recalled in a matching state (Godden & Baddeley, 1975; Smith & Vela, 2001). Whereas laboratory studies of emotion-state dependent recall have yielded mixed findings (Bower & Mayer, 1985; Eich, 1995), the dependency of new learning on emotional context is routinely observed in clinical settings (M. Craske et al., 2008). Patients with anxiety who form new associations between feared stimuli and neutral outcomes often find that when encountering these stimuli in new contexts, the associated anxiety reemerges (M. Craske et al., 2008). Here, we tested whether CMR3 can account for emotion-state dependent recall.

#### Simulation 3

##### Method

We simulated a delayed free recall task consisting of 60 lists of 10 neutral items. We simulated mood inductions (1) at the beginning of each encoding list and (2) immediately prior to recall. The order of mood inductions in each session was randomized, such that 30 lists had matching encoding and retrieval contexts (15 negative-negative, 15 positive-positive) and 30 lists had mis-matched encoding and retrieval contexts (15 negative-positive, 15 positive-negative). We then calculated the model-predicted recall probabilities and response times conditional upon matching negative, matching positive, mismatched negative-positive, and mismatched positive-negative, encoding and retrieval contexts. Finally, we ran this simulation with each set of parameters obtained by fitting CMR3 to individual subjects’ behavioral data in Simulation 1.

##### Results and Discussion

In this simulation, we tested whether virtual subjects recalled neutral items better when their emotional state during retrieval matched the emotional state that was present during encoding. The model-predicted recall probabilities for neutral items encoded and retrieved in matching emotional states were .67 (SEM = .027) for negative states and .69 (SEM = .025) for positive states. The recall probability for neutral items encoded in a negative state and retrieved in a positive state was 0.41 (SEM = .033), and the recall probability for neutral items encoded in a positive state and retrieved in a negative state was 0.38 (SEM = .035). This decrease in recall for neutral items in non-matching (vs. matching) emotional states during encoding and retrieval reflects the model’s core prediction: an item’s activation for retrieval depends on the similarity between the current context and the context associated with that item during encoding. In CMR3, emotional content is part of those contexts. Thus, the model can account for findings that new learning in clinical settings is enhanced when patients learn new material or associations in an emotional state that matches the emotional state in which they will later need to activate the new learning (M. Craske et al., 2008).

### Persistent Negative Mood

Negative mood that persists for two weeks or longer is a core feature of MDD. Stressful life events can trigger MDD, with the likelihood of a depressive episode increasing with the frequency of recent negative events (Kendler, Karkowski, & Prescott, 1998). Early developmental stressors, such as childhood abuse and neglect, also contribute to later risk of depression in adulthood (Jaffee et al., 2002; Kendler et al., 1998, 1995), perhaps in part by influencing how individuals recall their experiences and respond to new events. Going beyond modeling emotional effects on list memory, we seek to simulate an individual’s life history, and evaluate how prior and recent negative life events can lead to the development and persistence of negative mood through the mechanisms of our memory model.

### Simulation 4

Here we imagine human experience as unfolding in a sequence of events. We further consider each event as analogous to a list composed of sequentially experienced stimuli, each defined by their features, emotional valence, and associations to other stimuli. Mental life involves both experiencing these events (akin to learning a list) and thinking about them, or remembering them, at future moments (akin to recalling the list). These simplifying assumptions allow us to generalize the rich literature on list memory paradigms to humans’ experience at much longer time scales, albeit stripping away much of the complex structure of real-life events.

In this and subsequent simulations, we evaluate different assumptions regarding a simulated individual’s history with emotional experiences for that individual’s contemporaneous and subsequent mood. We then simulate the acquisition of these experiences using the learning rule embodied in CMR3 and the model’s assumptions regarding the evolution of context. We assume that individuals not only experience events, but also think about them, and simulate these internal reflections as a memory retrieval process akin to the free recall task. We model internal thinking as retrieval of stimuli, cued by the model’s continuously evolving representation of context.

These simulations address four key components of a theory of clinical disorder: how the disorder develops, what factors maintain it, the mechanisms of treatment, and what factors lead to sustained improvement versus relapse. These correspond to four distinct simulation epochs: A Developmental Period, a Neutral Period, a Treatment Period, and a Post-Treatment Period. Before describing how we simulated each epoch, we present methods for simulating individuals who vary in both the parameters that govern their memory processes and in their unique sequence of experiences.

#### Method

Using the model parameters obtained from our detailed fits to the behavioral data in Simulation 1, we created a sample of 97 virtual subjects. For each virtual subject, we simulated a life-course of events, each presented within a “list,” or sequence of stimuli, that comprises a larger episode in a virtual subject’s life. At the end of each such episode, or list, we allowed the model to engage in a moment of “free recall,” to model the spontaneous recall of these prior events, based on the literature that the same episodic memory system which supports recall of voluntary (intended) memories also supports recall of involuntary (spontaneous, unintended) memories (Berntsen, 2010). We simulated four periods of life events, each distinguished by their compositions of neutral, negative, and positive events: (1) a Developmental Period, (2) a Neutral Period, (3) a Treatment Period, and (4) a Post-Treatment Period.

The *Developmental Period* tests the model’s ability to predict different outcomes for virtual subjects depending on their history of early lifetime emotional events. During this period, virtual subjects experienced 50 sets, or “lists,” of 10 encoding events. We designed three such Developmental Periods for virtual subjects to experience. In the Low-Frequency condition, lists contained 40% neutral, 30% negative, and 30% positive events, resulting in a 1:1 ratio of negative-to-positive events. In the Moderate-Frequency condition, we increased the frequency of negative encoding events, such that lists contained 20% neutral, 60% negative, and 20% positive events, resulting in a 3:1 ratio of negative to positive events. In the High-Frequency condition, we further increased the frequency of negative events, such that lists consisted of 20% neutral, 70% negative, and 10% positive events, resulting in a 7:1 ratio of negative to positive events. Using a simulation approach allowed us to evaluate each condition (Low, Moderate, and High frequencies of negative events during the Developmental Period) for each virtual subject.

After the Developmental Period, virtual subjects experienced a *Neutral Period* in which they encoded and recalled 30 new lists, each consisting of 10 unique, neutral events. The Neutral Period served to isolate and identify the effects of prior learning during the three distinct Developmental Periods on subsequent mood and memory, without needing to filter out the effects of new emotional events.

A theory of emotional disorders must account not only for risk and maintaining factors, but also for which interventions reduce symptoms. After the Neutral Period, virtual subjects experienced a simulated *Treatment Period*. This phase served to evaluate whether CMR3 could predict the efficacy of positive event scheduling. In positive event scheduling– a core component of Behavioral Activation Therapy (BAT) for depression–patients engage with positive and meaningful activities (Beck et al., 1979; Jacobson et al., 1996; Lewinsohn, Sullivan, & Grosscap, 1980). Each simulated course of positive-event scheduling consisted of 20 new lists of 10 events. Each list had a heightened ratio of positive to negative events (3:1), having a list composition of 30% positive, 10% negative, and 60% neutral events.

Finally, after the Treatment Period, virtual subjects experienced a *Post-Treatment Period* evaluating their recovery. This phase served to evaluate the conditions under which CMR3 predicted relapse versus a sustained response to treatment. During the Post-Treatment Period, each virtual subject experienced 60 lists of predominantly neutral events intermixed with a small proportion of positive and negative events (80% neutral, 10% negative, 10% positive events).

##### Event Properties

For each event in the periods described above, we assigned the event’s properties of emotional valence (positive, negative, or neutral) and semantic relatedness to other events, as described below. For simplicity, we did not permit mixed-valence events. To simulate variability in the semantic associations among events, we drew the values for all inter-event similarities, regardless of events’ emotional valence, at random and without replacement from the same normal distribution (*µ* = 0.2, *σ* = 0.5). As a concrete example of these inter-item similarities, a birthday party and an engagement party might have a positive similarity of 0.8, whereas going swimming and filling out a tax form might have close to no semantic relationship, resulting in a score of -0.1 (Manning & Kahana, 2012; Manning, Sperling, Sharan, Rosenberg, & Kahana, 2012). We reviewed the resulting inter-item similarities to confirm that no draws had resulted in a perfect similarity score of 1.0. Thus, all simulated events were unique. We set *s*, which scales these inter-item similarity values and thus controls how much semantic associations guide memory retrieval, to a value of 0.15. This value allowed semantic mechanisms to contribute to the simulation without overshadowing the mechanisms of contextual episodic associations. Drawing inter-item similarity values for positive, negative, and neutral events from the same distribution ensured that differences in semantic associations could not explain the model-predicted mood and memory outcomes.

##### Simulated Mood and Recall

As each virtual subject experienced the simulated encoding events, we tracked the model-predicted levels of mood evoked by each event. To provide a concise graphical depiction of the model’s mood, we calculated a composite measure defined as the difference between the levels of positive and negative affect present in the context vector at that time.

During the session of free recall after each list, each virtual subject’s memory system spontaneously responds to the current context as a retrieval cue for encoded memories. We tracked the occurrences of *voluntary memories*, referring to memories from an intended (target) life-context. For simplicity, we operationalized the target context as the most-recent list of events. In addition, we tracked the occurrences of *involuntary memory intrusions*, referring to memories from an undesired and non-intended (non-target) life context. CMR3 models non-target life contexts as any of the lists preceding the most-recent list of events. For consistency with the list-learning literature, we refer to the former as “correct recalls” and the latter as “prior-list intrusions.” The *correct recall* metric refers to the probability of correctly recalling a target item, and it is calculated independently of the number of prior-list intrusions that occur in that list.

### Results

CMR3 predicted that different compositions of emotional events during a virtual subject’s early development influence their subsequent mood and memory. Using a simulation framework allowed us to disentangle the model-predicted, distinct contributions of the environment and the individual. Holding the parameters of each subject’s unique memory system constant, we ran the simulation multiple times for each subject, varying the emotional composition of their Developmental Period during each run.

In Figure 6, we illustrate the dynamics of emotion for one virtual subject, to demonstrate the simulation outputs in fine detail. The vertical axis tracks the model-predicted mood for this virtual subject. The graph’s height represents the positivity or negativity of model-predicted mood, calculated as the difference (subtraction) between the level of positive emotion and negative emotion present in the virtual subject’s emotional context at that point in the simulation. The color of each point on the graph represents the type of event, whether an encoding event or a retrieval event, that evoked the current mood state. Blue represents a negative event, pink a positive event, and gray a neutral event. In each simulation, a virtual subject encounters the same number of events.

**Figure 6.**
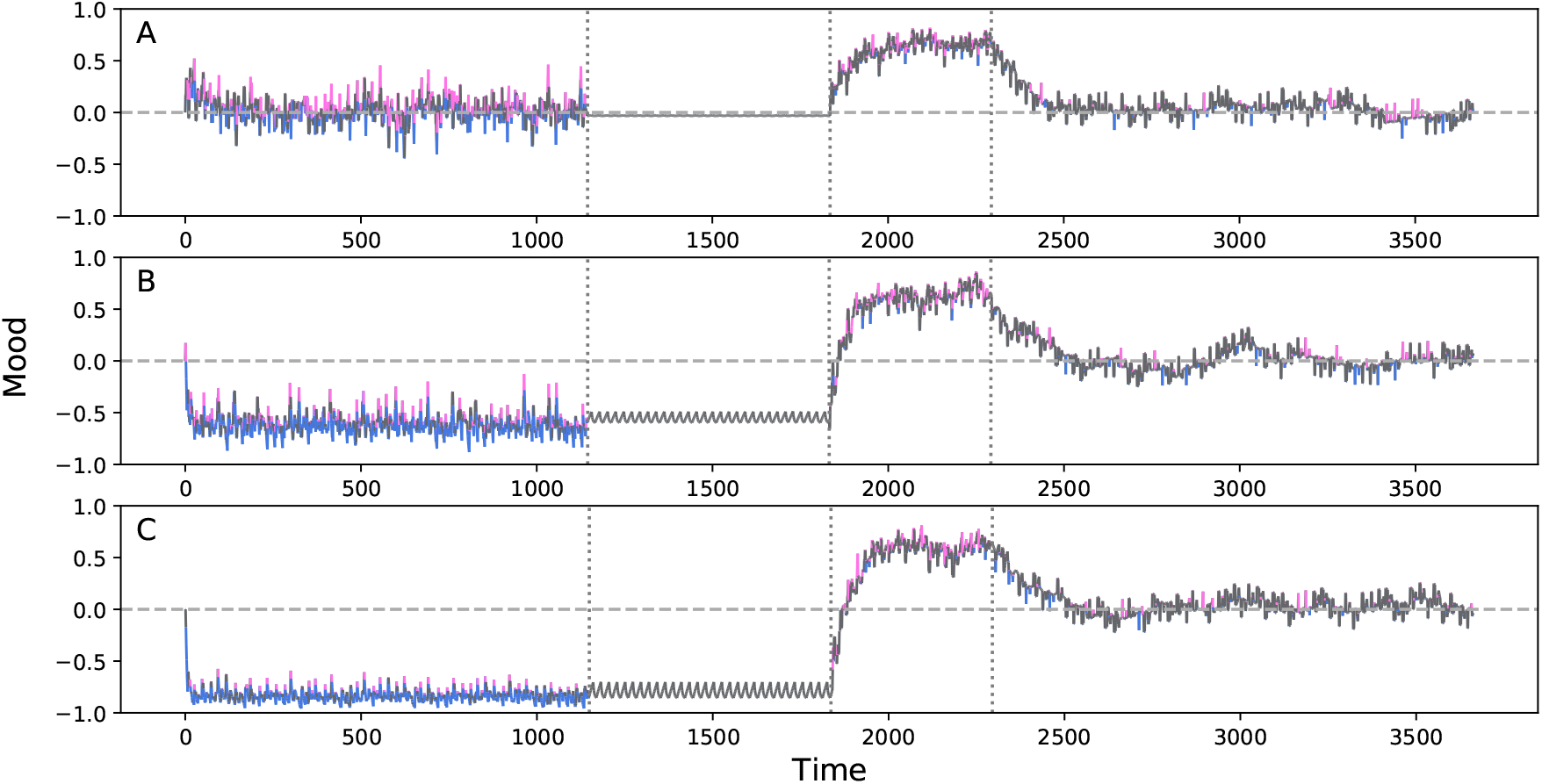
Frequency of Emotional Events Determines Evolution of Model-Simulated Mood. Across four simulation periods, delineated by vertical lines, one virtual subject experienced and remembered positive (pink), neutral (gray) and negative (blue) events. Vertical axis indicates model-simulated mood, ranging from -1 (max. negative) to +1 (max. positive). During an early Developmental Period, this virtual subject experienced varying proportions of positive and negative events, in each of three simulation runs. **A**. Equal proportions of negative and positive events (30% each). **B**. 60% negative and 20% positive events. **C**. 70% negative and 10% positive events. During a subsequent Neutral Period, virtual subjects experienced solely unique, neutral events. During a simulated course of positive event scheduling, virtual subjects experienced 10% negative and 30% positive events. During a final Post-Treatment Period, virtual subjects experienced 10% negative and 10% positive events.

The results demonstrate how the same individual would have responded to each of three different, early Developmental Periods (Figure 6): one having few negative events (panel A), one having a moderate number of negative events (panel B), and one having many negative events (panel C). The subsequent simulation periods (see *Method* for full details) demonstrate how a virtual subject may then develop negative mood, maintain it once prior negative events have abated (the Neutral Period), what factors contribute to recovery (the Treatment Period), and under what conditions the virtual subject relapses or sustains that recovery (the Post-Treatment Period). However, because recall is stochastic, the number of recalled memories can vary across simulations; therefore, the length of each simulation differs across panels.

Whereas figure 6 shows the dynamics of mood for an individual example subject, figure 7 displays the aggregated patterns of model-predicted mood and memory taken across all virtual subjects, averaged within each period of the simulation. CMR3 predicted an increase in negative mood with increasing frequency of early-life negative events (Figure 7, Panel A). Furthermore, the emotional context that accrued during the Developmental Period persisted into the Neutral Period, such that virtual subjects who experienced high rates of prior negative events developed persistent negative mood, even though they rarely recalled events from the Developmental Period itself (Figure 7, Panel B). Thus, the model predicted that negative mood can perpetuate from prior negative experiences even when a patient is not actively recalling them. Rather, recalling recent neutral events that had become associated to negative background context during their encoding, evoked the emotional context with which they had been encoded. Thus, the negative emotion from recent negative events generalized to neutral events, causing them to reactivate negative mood, even though these neutral events had only neutral properties in and of themselves.

**Figure 7.**
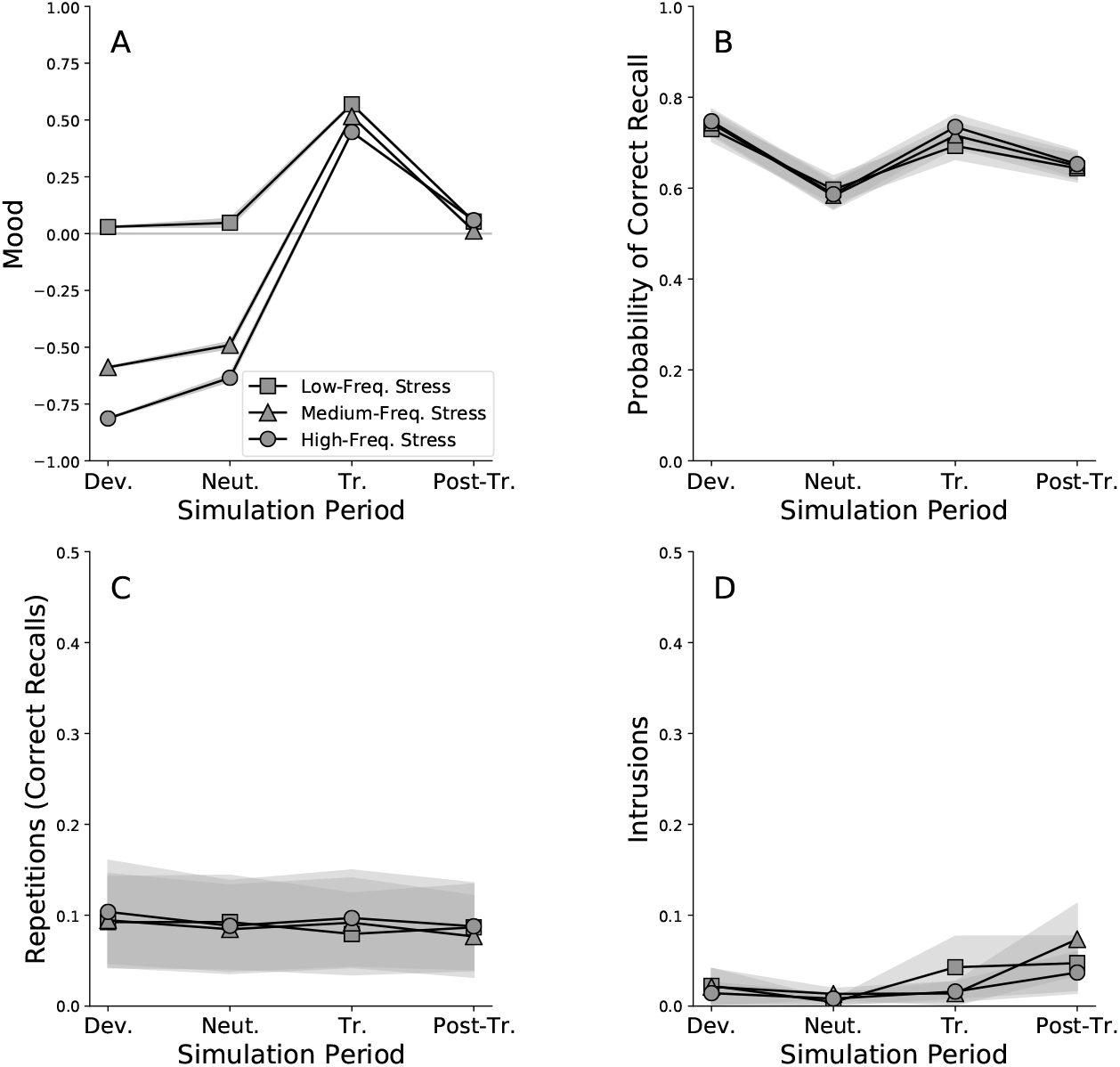
Aggregated Results for Negative-event Frequency Effects. Model-predicted patterns of mood and recall for Simulation 4, aggregated across all virtual subjects (i.e., parameter sets). Virtual subjects experienced four simulation stages: Developmental (Dev), Neutral (Neut), Treatment (Tr), and Post-Treatment (Post-Tr). During an early Developmental Period, virtual subjects experienced varying proportions of positive and negative events. Square markers represent an equal proportions of negative and positive events (30% each). Triangle markers represent a Developmental Period of 60% negative and 20% positive events. Circle markers represent a Developmental Period of 70% negative and 10% positive events. During a subsequent Neutral Period, virtual subjects experienced solely unique, neutral events. During a simulated course of positive event scheduling (“Treatment Period”), virtual subjects experienced 10% negative and 30% positive events. During a final Post-Treatment Period, virtual subjects experienced 10% negative and 10% positive events. **A**. Mean levels of virtual subjects’ mood. **B**. Probability of correct recall. **C**. Mean number of memory-repetitions per list. **D**. Mean number of memory-intrusions per list. Error regions represent ± 1 *S*.*E*.*M*., calculated across all parameter sets’ predictions.

Positive event scheduling produced an improvement in mood that persisted during the Post-Treatment Period, regardless of the frequency of early-life negative events (Figure 6 and Figure 7a). The persistence of negative mood in the Neutral Period prior to this intervention arose because, although neutral events in these simulations do not hold emotional content, neutral events will be bound to a variable emotional context. When bound to a relatively neutral or positive context, recalling such neutral events will wash out a strong negative context as time goes by, since remembering the event reactivates its associated, non-negative emotional context. Thus, neutral events can also attenuate the extremity of mood.

However, improved mood following a neutral event does not occur if the event has formed associations with negative contexts, as in the Moderate-Frequency and High-Frequency conditions. In this case, neutral events temporarily reduce negative mood when they occur, but when such neutral events are recalled, they will reactive their associated negative contexts, causing negative mood to persist. Overall, CMR3 predicts that both the valence of an event’s features and the valence of its associated emotional context combine to contribute to the current mood. Thus, encountering new neutral or positive events against the backdrop of a nonnegative emotional context is essential for mood to improve.

Figure 7b-d presents the average level of each type of recalled memory within each simulation period, averaged across virtual subjects. For consistency with list-learning paradigms, we use “correct recall” to refer to instances in which the virtual subject has recalled a memory that occurred within the intended (target) life-context (i.e., “list”). In our simulations, this corresponds to voluntary (strategic, or targeted) autobiographical recall (Berntsen, 2010). Also drawing from list-learning literature, we use the term “prior-list-intrusion” to refer to instances in which the virtual subject has spontaneously recalled an undesired memory from an unintended (non-target) life-context (i.e., a “prior list”). Because such memories intrude from a prior list, they are called *prior-list intrusions* (PLIs).

During the Neutral Period, the model predicted decreased probability of recall for recent neutral events while virtual subjects were still experiencing persistent low mood, due to the prior negative events (Figure 7b). Recall for these events improved during the simulated Treatment Period (Figure 7b). Prior theorists accounting for reduced recall of neutral or positive events during depressed mood states have proposed that such reduced recall arises from impairment of the neural structures that support the memory system (Austin, Mitchell, & Goodwin, 2001; Bremner, Vythilingam, Vermetten, Vaccarino, & Charney, 2004). Our results propose an alternative, or perhaps complementary, explanation. In a negative mood state, mood-congruent recall processes (see Simulation 2) should decrease recall for neutral and positive stimuli. Since cognitive tests often use neutral stimuli, our simulations suggest the importance of future research to test whether findings of reduced memory in depressed patients may reflect the mismatch between items’ emotional properties and the subject’s emotional state, rather than (or perhaps as a compounding factor to) memory system impairment.

### Repetition Effects

Repeating an event increases its subsequent memorability, especially when the repetitions occur at distributed points in time (Lohnas & Kahana, 2014). Repeated exposure to severe, traumatic stressors correlates with heightened depressive symptoms (Herman, 1992) and more varied and severe PTSD symptoms (Briere, Kaltman, & Green, 2008; Cloitre et al., 2009). Even mild stressors, when repeated, can lead to depression-like effects in animal models, including learned helplessness and decreased pleasure-seeking behaviors (Cabib & Puglisi-Allegra, 1996; Willner, 1997). In human adults, daily hassles correlate with the persistence of clinically significant distress (Depue & Monroe, 1986).

Simulation 5 tests CMR3’s ability to capture the classic finding that low-arousal, but chronic negative events correlate with negative mood (Depue & Monroe, 1986). CMR3 generates a novel prediction: namely, that chronic, low-arousal negative events perpetuate negative mood via the mechanism of spaced repetitions, which heighten the memorability of negative events and, in the process, crowd out the activation of positive memories during retrieval.

### Simulation 5

#### Method

Virtual subjects experienced a Developmental Period, a Neutral Period, a Treatment Period, and a Post-Treatment Period. The design and event composition of each period was identical to Simulation 4. Virtual subjects experienced either low-, moderate-, or high-frequency negative events during the Developmental Period. However, during this period, one negative event repeated, occurring once every ten lists. This repetition schedule allowed us to evaluate the effects of repetition without substantially increasing the frequency of negative events during the Developmental Period.

#### Results

Memory of the repeated negative-event intruded into the virtual subjects’ memories during the Neutral Period, even though neither this repeated negative-event nor any other negative events had occurred during that period. In comparison, during Simulation 4, in which all negative events were unique, no memories of negative events intruded during the Neutral Period. Thus, spaced repetitions predisposed this negative event to spontaneously reactivate during recall periods. Two distinct groups emerged among the virtual subjects. One group experienced a high frequency of negative-memory intrusions, operationalized as one or more intrusions per list (High-Intrusions Group, N = 12). The second group exhibited almost zero memory-intrusions of the negative event (Low-Intrusions Group, N = 85) and more positive mood. Thus, different memory parameters predisposed some virtual subjects to experience heightened vulnerability to intrusive memories of negative events and corresponding negative mood. Figure 8 shows the effects of repetition on mood and memory within each group.

**Figure 8.**
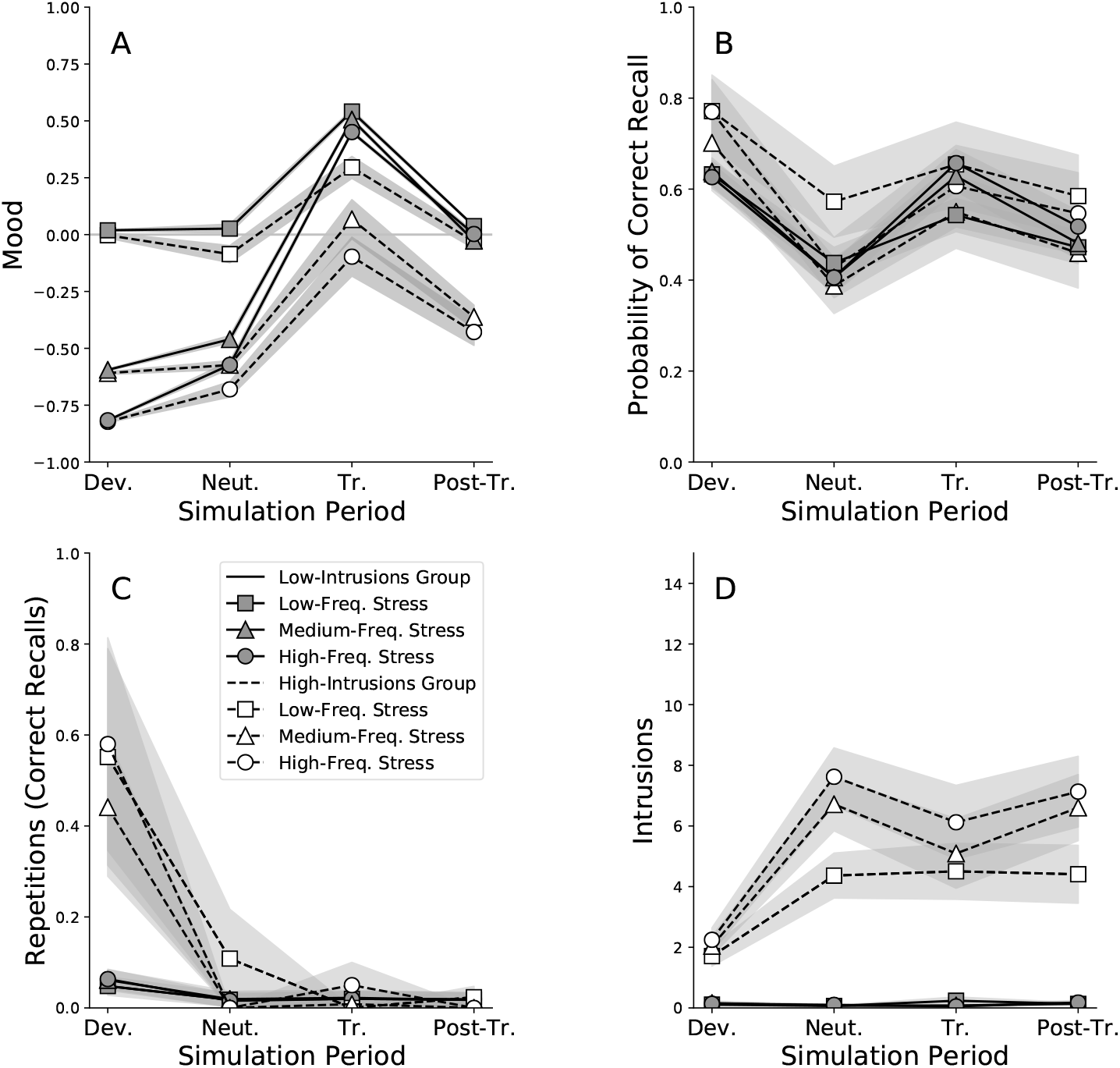
Effects of a Repeated Negative Event on Mood and Memory. Each panel shows results for Simulation 5, averaged across simulated subjects, in the Developmental (Dev), Neutral (Neut), Treatment (Tr), and Post-Treatment (Post-Tr) periods. Virtual subjects experienced either low, moderate, or high-frequency negative events during the Developmental Period, and one negative event repeated every 10 lists. All virtual subjects then experienced a period of neutral events, a simulated period of positive event scheduling (“Treatment Period”), and a Post-Treatment Period of predominantly neutral events. **A**. Mean levels of virtual subjects’ mood. **B**. Probability of correct recall. **C**. Mean number of memory-repetitions per list. **D**. Mean number of memory-intrusions per list. Solid lines show virtual subjects with low memory-intrusion counts (Low-Intrusions Group). Dashed lines show virtual subjects with high memory-intrusion counts (High-Intrusions Group). Error regions represent ± 1 SEM, calculated across virtual subjects.

Repeated negative events also affected the ability of positive event scheduling to lift depressed mood. Examining the simulation runs for individual virtual subjects revealed that heightened rates of positive events during the Treatment Period temporarily lifted mood. However, when intrusive memories of negative events occurred during recall, they continued to evoke negative context. Further, the heightened activation of negative emotional context reduced recall of more recent positive events, due to the process of mood-congruent recall (see Simulation 2), and as observed in real depressed participants (R. E. Nelson & Craighead, 1977). For the High-Intrusions group, these combined effects worsened mood during the Treatment and Post-Treatment periods relative to the Low-Intrusions group (Figure 8.

Simulation 5 proposes a novel mechanism by which such chronic, yet mild negative events may contribute to persistent negative mood. In retrieved-context theory, repetition associates the negative event with a wider variety of contexts. This increases the potential retrieval cues that can prompt recalling the negative event, which both raises the probability of recalling the negative event and potentiates it for activation in contexts where it does not belong (i.e., as a memory intrusion). Compounding this effect, the principles of mood-congruent recall predict worsened mood for two reasons: reactivating negative context increases the probability of recalling subsequent negative memories, and decreases the probability of recalling positive memories that might otherwise lift mood (see Simulation 2).

Our results highlight the importance of directly targeting intrusive memories of negative events in treatments for depression. Indeed, treatments that directly target intrusive memories in depression have shown promising results (C. R. Brewin et al., 2009), as have treatments targeting perseveration about distressing memories in the context of rumination (E. R. Watkins et al., 2011). Next, we examine the model’s ability to characterize the mechanisms of intrusive, non-voluntary memories of high-arousal negative events.

### Intrusive Memories

In the aftermath of extreme stress, PTSD patients report that involuntary, intrusive, and often repetitive, memories of the traumatic event constitute their most distressing symptom (Ehlers & Steil, 1995). For intrusive memories in PTSD, stimuli that resemble those directly preceding the traumatic event serve as a strong cue of memory intrusions (Ehlers et al., 2004). Such stimuli, here called “trauma cues,” often share little or no semantic similarity to the event itself (Ehlers & Clark, 2000). For example, a patient who was assaulted near a water fountain may experience flashbacks when hearing the sound of running water, even though there is no semantic association between water and assault. The tendency of contextual cues, including perceptual, temporal, and spatial elements of context, to activate trauma-memory intrusions is well known in PTSD. However, intrusive memories of painful events are found in numerous disorders beyond PTSD (C. R. Brewin, Gregory, Lipton, & Burgess, 2010; Hackmann et al., 2000; Muse et al., 2010) and often involve stressors not traditionally considered to be traumatic, such as social rejection or perceived failure (A. D. Williams & Moulds, 2007).

Leading treatments for PTSD, such as prolonged exposure therapy (Foa & Kozak, 1986), target trauma-memory intrusions through two routes. During *in vivo* exposures, patients repeatedly approach the contextual cues that trigger intrusions of the trauma memory. During *imaginal exposure*, patients repeatedly revisit the trauma-memory itself. Simulation 6 asks whether CMR3 can account for the finding that neutral stimuli preceding a negative event can reactivate intrusive memories of that event. Further, this simulation examines the effects of avoiding such cues, versus repeatedly approaching these cues during *in vivo* exposures.

### Simulation 6

#### Method

To model trauma-memory intrusions, Simulation 6 uses the *externalized free recall* (EFR) paradigm (Zaromb et al., 2006; Kahana et al., 2005; Lohnas et al., 2015). In EFR, subjects study lists of items and then report all items that enter their mind during the recall period, whether or not they intended to recall them. Thus, subjects report both voluntary (strategic) and involuntary (spontaneous) recalls. In autobiographical memory, although subjects may experience voluntary and involuntary recall as distinct phenomena, literature suggests that both processes share and operate through similar mechanisms (Berntsen, 2010; Kahana et al., 2005). Thus, we operationalized memory-intrusions as spontaneous memories that a virtual subject recalls from an unintended and undesired prior life-context, as defined by a “list of items” in our simulation. We thus refer to such responses as *prior-list intrusions*, or PLIs.

During EFR, subjects exhibit varying degrees of control over, and awareness of, PLIs (Zaromb et al., 2006), analogous to spontaneous trauma memories in PTSD. For example, in PTSD, patients often report experiencing trauma-memory intrusions spontaneously, or “out of the blue.” However, further investigation typically reveals that the memory was in fact cued by a temporal, spatial, or other contextual cue associated with the trauma (Ehlers & Clark, 2000). Even healthy individuals spontaneously recall neutral memories that are unintended and undesired, highly vivid, and which they may not recognize as being an intrusion (Berntsen & Rubin, 2002; Berntsen, 2010; Staugaard & Berntsen, 2014; Zaromb et al., 2006; Kahana et al., 2005; Roediger & McDermott, 1995).

Simulation 6 followed the same format as Simulation 4 but differed in several crucial ways. First, all virtual subjects experienced the same Developmental Period of 50 lists of 10 encoding events, each consisting of 40% neutral, 30% positive, and 30% negative events. Within this Developmental Period, we selected a single, random, negative item to serve as a traumatic event. We set the arousal parameter, *ϕ*_*emot*_ equal to 5.0 to amplify the strength of association between the traumatic event and its temporal context during encoding (see Equation 8). We modeled trauma-associated stimuli, or “trauma cues,” as the two neutral stimuli that had directly preceded the traumatic event during encoding.

Then, the simulation tested CMR3’s ability to capture findings that avoiding trauma cues perpetuates trauma-memory intrusions, whereas repeatedly approaching trauma cues, such as during *in vivo* exposures, alleviates reexperiencing symptoms (Foa, Hembree, & Rothbaum, 2007). After the Developmental Period, each virtual subject experienced one of three intervention conditions. In the *Avoidance Condition*, virtual subjects avoided (i.e., never reencountered) all trauma cues, thus never reencoding them in new contexts. In the *Single Exposure* condition, virtual subjects approached (i.e., reencountered) and thus reencoded their trauma cues a single time, but then returned to avoiding their trauma cues afterward. In the *Repeated Exposure* condition, virtual subjects approached, and thus reencoded, their trauma cues eight times, corresponding to the typical number of weeks that patients are instructed to engage in *in vivo* exposure exercises during a course of prolonged exposure therapy (Foa et al., 2007). The Avoidance, Single Exposure, and Repeated Exposure conditions allowed us to test the model’s ability to capture a dose-response relationship between the number of times a virtual patient approached versus avoided trauma cues.

Simulation 6 modeled avoidance or approach of trauma cues by embedding these stimuli as new encoding events during each intervention condition, as follows. Each intervention condition began with 10 lists of unique, neutral events, followed by 100 lists of the given intervention. In the Avoidance condition, these 100 lists consisted solely of new, unique neutral events. In the Single Exposure condition, these 100 lists consisted of new, unique neutral events, and after 10 lists, we embedded the two neutral trauma cues in one list. In the Repeated Exposure condition, these 100 lists consisted of new, unique neutral events, and we embedded the two neutral trauma every 10th list, at which point the virtual subjects reencountered the trauma cues and reencoded them.

### Results

Simulation 6 evaluated whether CMR3 could predict the effects of “trauma cues” on cueing intrusive involuntary recall of that event (Ehlers & Clark, 2000). Figure 9 shows the rate of trauma-memory intrusions for each virtual subject during the Neutral Period that followed exposure to the traumatic event. CMR3 predicted that all subjects would spontaneously recall the traumatic event for a short time after exposure. However, by the Neutral Period, two groups emerged. For one group, comprising 85 of the 97 virtual subjects, trauma-memory intrusions had subsided to fewer than one intrusion per recall list by the start of the Neutral Period (Low-Intrusions Group). In the second group, 12 virtual subjects continued to experience trauma-memory intrusions at least once per recall list by the start of the neutral-events only period (High-Intrusions Group). This resembles actual rates of chronic PTSD after trauma exposure, which typically range from 5-10% but can be higher depending on the type and severity of the trauma (Bonanno, Westphal, & Mancini, 2011).

**Figure 9.**
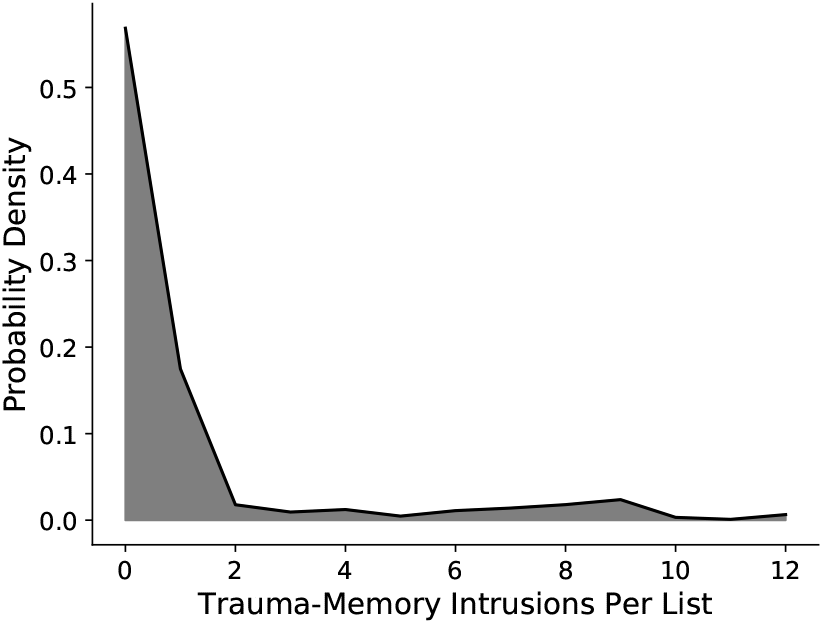
Distribution of Trauma-memory Intrusions per List for Virtual Subjects. Whereas 85 of 97 virtual subjects exhibited a mean of fewer than one trauma-memory intrusion per list during the period of neutral events following their trauma, 12 virtual subjects experienced trauma-memory intrusions at higher frequencies.

We assessed the model-predicted effects of encounters with trauma cues, defined as the two neutral stimuli that had directly preceded the traumatic event during encoding. We evaluated these predictions in each of three conditions: In the Avoidance condition, virtual subjects never approached trauma cues; in the Single-Exposure condition, virtual subjects approached trauma cues once; and in the Multiple-Exposures condition, virtual subjects approached trauma cues eight times. Figure 10 shows the model-predicted frequency of trauma memory intrusions (upper panels) and the model-predicted mood (lower panels) separately for the High-Intrusion Group (left panels) and the Low-Intrusion Group (right panels).

**Figure 10.**
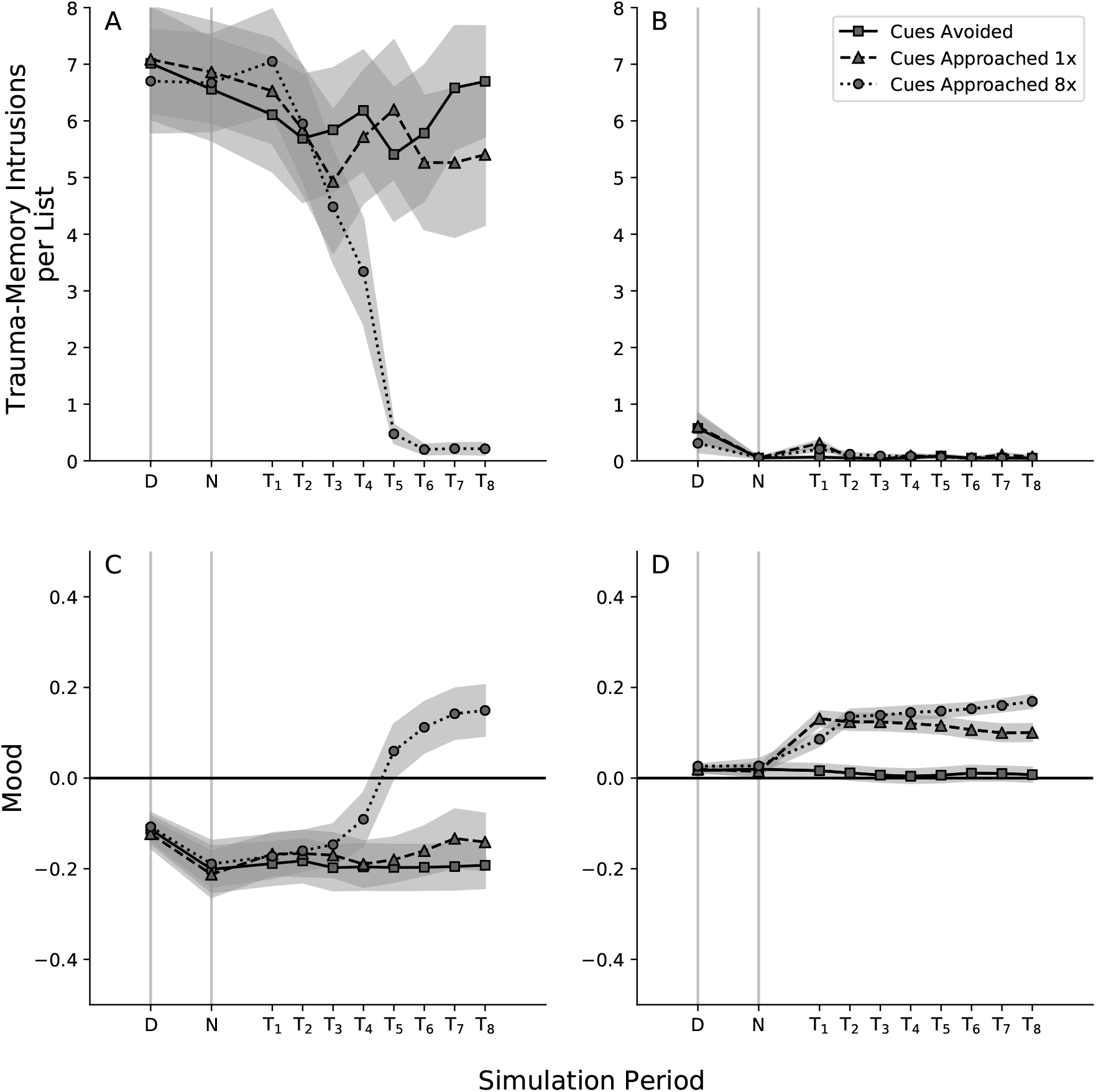
Model-predicted mood and trauma-memory intrusions after a traumatic event. Results are aggregated within two groups of virtual subjects: those with minimal or no trauma-memory intrusions after the traumatic event (Low-Intrusions Group) and those with frequent trauma-memory intrusions (High-Intrusions Group). Virtual subjects experienced a Developmental Period containing one high-arousal, negative event (traumatic event), followed by one of three intervention conditions. Square markers represent a baseline condition, in which the Neutral Period consists solely of unique, neutral events (avoidance condition). Triangle markers represent a single exposure to two neutral trauma cues (single exposure condition). Circle markers represent eight exposures to the same two neutral trauma cues (repeated exposures condition). **A**. Trauma-memory intrusions over time, High-Intrusions Group. **B**. Trauma-memory intrusions over time, Low-Intrusions Group. **C**. Mood over time, High-Intrusions Group. **D**. Mood over time, Low-Intrusions Group. D = Developmental Period following the traumatic event. N = The Neutral Period prior to the potential re-encounters with trauma cues. T_1−8_ = each of the eight time-points at which a virtual subject has an opportunity to either approach or avoid trauma cues.

Virtual subjects in the Low-Intrusions group experienced trauma-memory intrusions for a short period following the traumatic event, after which their frequency of trauma-memory intrusions returned to zero or near-zero levels (Figure 10b). Re-encountering trauma-associated stimuli a single time cued a temporary increase in memory-intrusions of the event for these subjects, which then also returned to near-zero levels (Figure 10b). This result is consistent with findings that the temporal context of a traumatic event can serve as powerful cues for flashbacks in PTSD, even when these contextual stimuli are neutral and semantically unrelated to the event (Ehlers et al., 2004). Retrieved-context theory predicts this finding for the stimuli that precede rather than follow a traumatic event due to the *temporal contiguity effect*. As shown in Figure 2C, recalling one item cues recall of other, temporally-associated items in a forward-biased manner (Healey et al., 2019). This forward-bias arises because encountering an item or event causes a persistent change in context (Equation 1), such that the item’s retrieved context becomes a stronger cue for the items that followed it during encoding, rather than those that preceded it.

Virtual subjects in the High-Intrusions group experienced frequent trauma-memory intrusions that persisted into the Neutral Period. Avoiding trauma cues (Avoidance condition), or encountering them only once (Single-exposure condition), allowed trauma-memory intrusions to persist at high levels (Figure 10a). In these intervention conditions, mood for virtual subjects in the High-Intrusions group did not improve. However, repeatedly approaching the trauma cues reduced trauma-memory intrusions to a frequency comparable with that of the Low-Intrusions Group, i.e., nearly zero (Figure 10a). As virtual subjects reencountered the trauma cues, they associated these cues with more varied, non-traumatic contexts. As a result, the cues became associated with, and began to evoke, a wider variety of recall outputs, thereby decreasing their tendency to cue recall of the traumatic event.

Repeatedly encountering the trauma cues improved mood for virtual subjects in both groups (Figure 10c-d). However, for the High-Intrusions group, mood only improved once repeated exposures reduced the frequency of distressing trauma-memory intrusions (Figure 10c). Then, the trauma cues began to evoke prior positive contexts that they had been associated with during the Developmental Period, prior to the traumatic event. An important nuance is that the trauma cues themselves were neutral, not positive stimuli, and no positive stimuli occurred during the intervention phase of the simulation. Rather, prior to becoming associated with the traumatic event, the neutral trauma cues had been associated with a mildly positive context evoked by an earlier, positive stimulus. Reencountering, and thus associating the trauma cues to new contexts, decoupled them from the traumatic event, decreased their tendency to reactivate the trauma-memory, and restored their tendency to evoke their prior, positive associations and new, non-traumatic associations.

Thus, CMR3 proposes a novel account of how, by restructuring episodic association networks in memory, trauma survivors can not only decrease negative affect resulting from trauma-associated stimuli, but also, recover prior enjoyment of such stimuli. For example, in exposure-based treatments, survivors of intimate partner violence not only can habituate to feared (but safe) romantic interactions, but also can recover feelings of joy and affection in romantic relationships.

For addressing both trauma-memory intrusions and negative mood, CMR3 predicted symptom change as the result of associating new neutral or positive contexts to the trauma cues, rather than replacing old ones. Simulation 6 provides independent corroboration of emerging findings that, even after successful exposure treatment, patients’ fear responses can reemerge when they encounter previously-feared stimuli in contexts where new learning did not take place (M. Craske et al., 2008). Retrieved-context theory proposes that the old responses associated with the distressing stimulus are still latent and can later reemerge under two conditions. Either the patient might encounter a contextual cue that is more strongly associated with the old responses than with new responses, or the current context might contain few elements of the context in which the new learning took place. This pattern is consistent with principles of inhibitory learning (M. Craske et al., 2008). However, using the framework of episodic memory processes, via restructuring the network of memory-to-context associations, enables our model to predict the reemergence of trauma-relevant emotions beyond fear, such as guilt, shame, sadness, or anger, when a trauma-memory is reactivated by a context in which new learning has not taken place.

CMR3 captured the tendency of stimuli preceding the traumatic event to serve as a potent cue for trauma-memory intrusions, such as the flashbacks characteristic of PTSD. The model predicted that *in vivo* exposures in Prolonged Exposure Therapy (PE; Foa & Kozak, 1986) restructure context-to-memory associations in episodic memory, and thereby reduce patients’ rate of trauma-memory intrusions. Simulation 7 models the effects of individual differences, such as emotion dysregulation, on the development of trauma-memory intrusions and negative mood.

### Emotion Dysregulation

#### Motivation

Difficulties understanding and regulating the intensity and valence of one’s emotional experiences increases the risk of developing PTSD and can lead to more severe responses to trauma cues (Forbes et al., 2020; Tull, Barrett, McMillan, & Roemer, 2007; Tull, Berghoff, Wheeless, Cohen, & Gratz, 2018; Messman-Moore, Walsh, & DiLillo, 2010). Simulation 7 examines how individual differences in emotion dysregulation moderate the occurrence of, and emotional response to, intrusive memories. Emotion dysregulation is a multifaceted construct (Gratz & Roemer, 2004), including not only the intensity of responding to emotional events (see Simulations 6, 8, and 9), but also how quickly a person incorporates the content of new emotional experiences and releases the effects of prior ones. Here, we tested CMR3’s predictions for the effects of quick versus slow incorporation of new emotional context on the frequency of trauma-memory intrusions and the level of mood.

### Simulation 7

#### Method

As in Simulation 6, virtual subjects experienced an early Developmental Period including a mixture of emotional and neutral events, followed by a Neutral Period of solely neutral events (see Simulation 6, “Avoidance Condition”). Using the 97 sets of parameters in the prior simulations, we modified the *β*_*emot*_ parameter to create two new groups of virtual subjects: a Slow Emotional-Updating Group (*βemot* = .3, N = 97 subjects) and a Fast Emotional-Updating Group (*β*_*emot*_ = .8, N = 97 subjects). To isolate the specific effects of changing the rate of emotional context updating, we varied only *β*_*emot*_ across groups. All other parameters remained fixed cross virtual subjects, and matched to those used in Simulation 6. All other methods carried over from Simulation 6.

#### Results

Virtual subjects in the Fast Emotional-Updating Group exhibited worse mood during the neutral period (M = -.08, SE = .02) compared to those in the Slow Emotional-Updating Group (M = -.02, SE = .01), *t*(192) = 2.15, *p* = .03. The frequency of trauma-memory intrusions did not reliably differ across groups (*M*_*f ast*_ = 0.98, *M*_*slow*_ = 0.94, *t*(192) = -0.10, n.s.).

Our results reflect that emotion dysregulation is multifaceted (Gratz & Roemer, 2004). Simulations 6, 8, and 9 demonstrate CMR3’s prediction that heightened intensity of emotional responding, one component of emotion dysregulation, reliably increases the rate of trauma-memory intrusions and decreases mood. The current simulation revealed that the rate of emotional context updating worsens mood but does not affect the frequency of trauma-memory intrusions. Across these two dimensions of emotion dysregulation, CMR3 provides a novel framework for identifying how individual differences predispose a given subject to psychopathology, by evaluating how distinct parameter values affect memory and mood.

### Intrusive Memories: Vividness and Nowness

#### Motivation

Vivid, unwanted and intrusive memories of painful events are best known in PTSD but occur widely across emotional disorders (C. Brewin, 2010), including depression (Reynolds & Brewin, 1999), social phobia (Hackmann et al., 2000), obsessive compulsive disorder (Speckens et al., 2007), health anxiety (Muse et al., 2010), body dysmorphic disorder (Osman et al., 2004), and agoraphobia (Day et al., 2004). Furthermore, in the flashbacks characteristic of PTSD, intrusions can evoke a sense of “nowness,” or a feeling that the memory is happening again in the here-and-now. Simulation 7 asks whether our model’s proposed mechanism for generating trauma-memory intrusions (heightened arousal during encoding), also predisposes intrusions to to be recalled more vividly and with a heightened sense of “nowness.”

### Simulation 8

#### Method

As in Simulation 6, virtual subjects experienced an early Developmental Period with 50 lists of 10 stimuli, each comprising 30% negative, 30% positive, and 40% neutral events. One item served as a traumatic event, which we embedded in the middle of its list (serial position 6) to minimize primacy and recency effects. The parameter *ϕ*_*emot*_, which scales the strength of context-to-item associations during encoding, operationalized negative arousal during the traumatic event. Varying *ϕ*_*emot*_ from 2.0 to 5.0 tested how negative arousal affects the vividness and nowness of subsequent trauma-memory intrusions. After the Developmental Period, virtual subjects experienced a Neutral Period containing 110 lists of new, neutral events, during which we examined the vividness and “nowness” properties of any trauma-memory intrusions that occurred.

CMR3 operationalizes “nowness” as the dot-product similarity between the subject’s current context and the context reactivated by the trauma-memory during recall, *c*_*sim*_ = **c**^*IN*^ · **c**_*t*_). If the similarity between encoding and retrieval contexts exceeds *c*_*thresh*_, then the subject experiences the memory as belonging to the most recent “now” context. *c*_*thresh*_ thus governs a subject’s ability to filter out unwanted memory intrusions and other extraneous mental content. A higher *c*_*thresh*_ value corresponds to a greater degree of cognitive control for a particular individual. CMR3 operationalizes vividness as how strongly context reactivates a memory’s features during recall. Mathematically, this corresponds to the item activation values that result from cueing the Hebbian weight matrices with the current recall context, **a** = *M*^*CF*^**c**_*R*_ (Equation 10).

#### Results

Heightened negative arousal during the traumatic event increased both the vividness and nowness of subsequent trauma-memory intrusions (Figure 11). In CMR3, this occurs for three reasons. First, negative arousal strengthens associations between the memory and its context during encoding, resulting in heightened reactivation of memory features during recall, and thus heightened vividness. Second, heightened negative arousal potentiates the memory system’s associative networks, such that trauma-memories require lower levels of associated contextual cues to reactivate them, and the trauma-memory begins to intrude more frequently.

**Figure 11.**
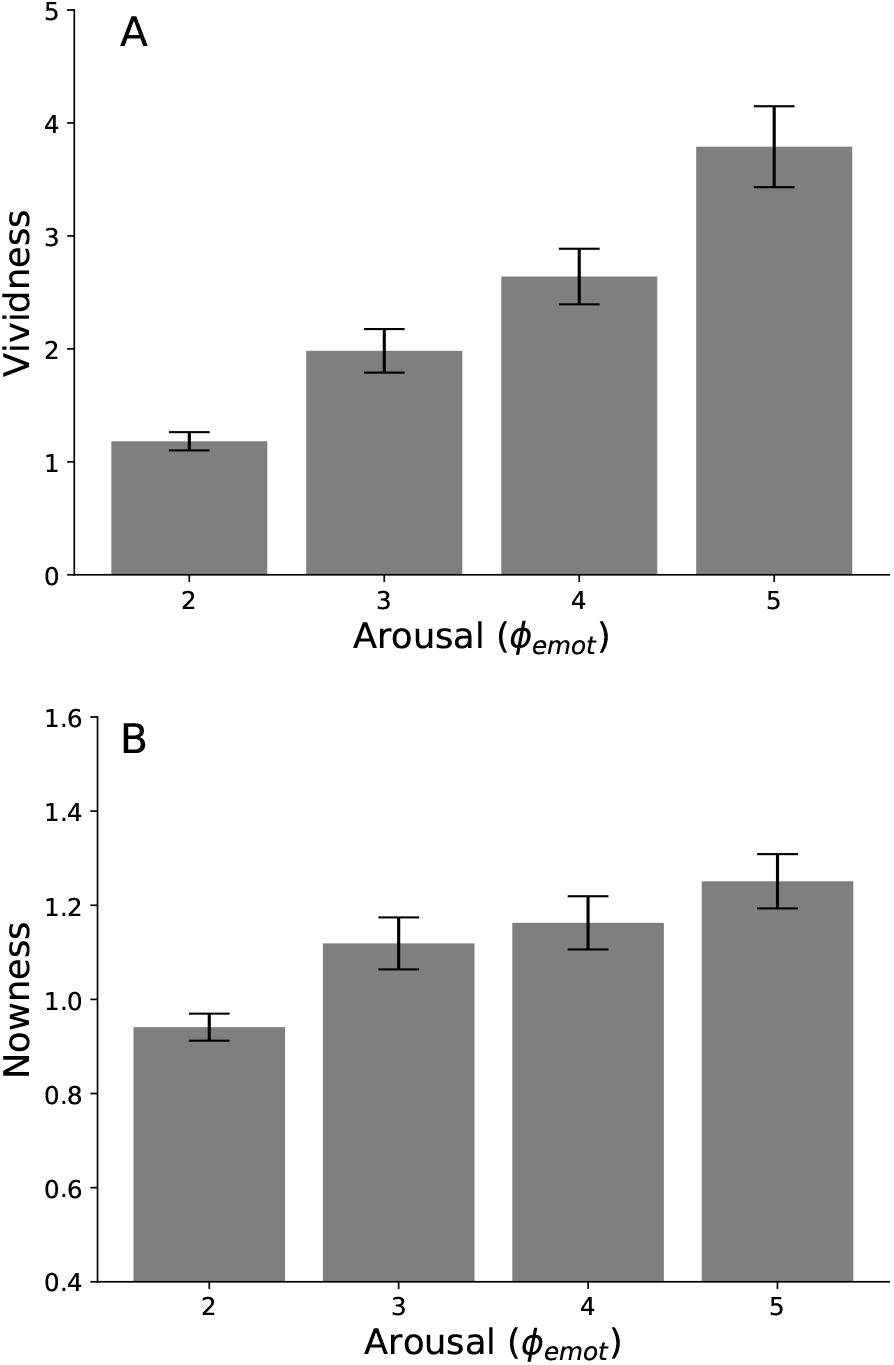
Negative Arousal Increases Model-Predicted Vividness and Nowness of Memories. **A**. Vividness (activation of a retrieved-memory’s features) as a function of negative arousal present during encoding (*φ*_*emot*_). **B**. Nowness (similarity match between encoding and retrieval contexts) as a function of negative arousal present during encoding (*φ*_*emot*_). Error regions represent ± *S*.*E*.*M*. across virtual subjects.

In addition, patients might introduce and repeatedly refresh the presence of trauma-associated context by ruminating about the trauma (Michael, Halligan, Clark, & Ehlers, 2007). Thus, subjects’ new contexts already contain traumatic content, resulting in a stronger match with the newly reactivated trauma context. Third, because traumatic events evoke strong negative emotional context, recalling a traumatic memory flavors the current context with negative features. The current context becomes more negative; thus, the emotional context present just prior to retrieval is a stronger match for the negative emotional context of the memory. The more-negative current context becomes more likely to cue the reactivation of further negative memory-intrusions, not necessarily limited to trauma-memory intrusions, due to mood-congruent recall (See Simulation 2). In turn, such memories flavor context with their own negative features, creating a snowball effect.

Because heightened negative arousal potentiates the associations between contextual cues and the trauma memory (Talmi et al., 2019), reduced levels of trauma-context are needed to cue spontaneous, involuntary recall of the trauma-memory. When low levels of trauma-associated context are present, one might predict a low match between the current context and the context reactivated by the trauma-memory, resulting in a reduced “nowness” value. However, even when external stimuli contain low levels of trauma-related perceptual features or spatiotemporal context, ruminating on the trauma memory will maintain internal representation of trauma-related context. In addition, one of the strengths of the eCMR and CMR3 models is the ability to account for a strong overlap in the source-memory components of encoding and retrieval contexts, such as emotional states, even when overlap in temporal contexts is reduced. Since patients with PTSD often experience a persistent sense of threat and other negative emotions (American Psychiatric Association, 2013), the retrieval context will be a stronger match for the negative emotional context that a trauma-memory reactivates, thus heightening the degree of contextual overlap and the resulting vividness or nowness.

### Prolonged Exposure Therapy

#### Motivation

Prolonged exposure therapy (PE; Foa & Kozak, 1986) is a highly effective treatment for trauma-memory intrusions and other symptoms in PTSD (Powers, Halpern, Ferenschak, Gillihan, & Foa, 2010; Tran & Gregor, 2016). PE has two core components. During *in vivo* exposure exercises, patients systematically approach feared stimuli that they have been avoiding due to their associations with the traumatic event (see Simulation 6). During *imaginal exposure* exercises, patients intentionally reactivate the traumatic memory and imagine it as though it is happening again in the here-and-now. Here, we test the ability of CMR3 to account for the efficacy of PE via the mechanism of restructuring episodic context associations. Further, we test CMR3’s ability to account for a pattern of reduced negative emotion in response to trauma memories that occurs across, rather than within, treatment sessions (M. Craske et al., 2008; Jaycox, Foa, & Morral, 1998).

### Simulation 9

#### Method

Simulation 6 demonstrated how repeated exposures to feared cues, and contexts associated with a traumatic event, gradually reassociates those cues to new, safe stimuli. In PTSD, the trauma memory is itself a feared stimulus that patients with PTSD avoid. Among other benefits, imaginal exposure decreases patients’ distress while reexperiencing the trauma memory, both in cases in which the patient recalls it voluntarily or in which it intrudes spontaneously. Simulation 9 tests CMR3’s ability to account for this phenomenon. In line with Foa et al. (2007), we modeled imaginal exposure as a process of reactivating the trauma memory, fully experiencing the memory and its emotional content as though it is happening again in the present. Then, we asked whether CMR3 can capture classic patterns of symptom improvement in PE via the mechanism of reencoding the memory in association with a new, safe and supportive context (e.g. the therapy office).

As in Simulation 6 (Intrusive Memories), virtual subjects experienced a Developmental Period consisting of 50 lists of 10 encoding events. Lists comprised the same proportion of positive, negative, and neutral items as in Simulation 6. One negative event during the Developmental Period served as the traumatic event. Here, however, we selected a very high value of *ϕ*_*emot*_ = 50.0, significantly amplifying the strength of the encoded associations between the traumatic event and its context. This high value ensured that all subjects would experience trauma-memory intrusions in the subsequent Neutral Period, and it served as a strong test of CMR3-predicted efficacy of imaginal exposure in reducing the activation of distress during trauma-memory intrusions.

After the Developmental Period, virtual subjects experienced a Neutral Period consisting of 110 lists of neutral events. In the Neutral Period, virtual subjects first experienced 10 lists of new, neutral items, followed by one of four interventions: (1) a Baseline condition, (2) an Imaginal Exposure condition, (3) a Positive Events control condition, and (4) a Re-encoding control condition. Due to the simulation framework, the same virtual subjects could experience each intervention condition on new runs of the simulation, allowing us to isolate the effects of the environment (the specific stimuli encountered) versus individual differences (defined by subjects’ parameter sets, held consistent across simulations).

During the Baseline condition, the Neutral Period consisted solely of new, neutral events. The Baseline condition served as a manipulation check to ensure that the chosen level of arousal resulted in high rates of trauma-memory intrusions for all virtual subjects. The Imaginal Exposure condition simulated the re-encoding of traumatic memories in safe, therapeutic contexts by re-introducing the trauma-memory as a new encoding event during the Neutral Period at five regular intervals, each spaced 20 lists apart. Prior to the re-encoding of the trauma memory, we modeled a safe context by introducing a positively valent stimulus. To reduce the risk of “overly positive” results in the Imaginal Exposure condition, these positive events had the same low level of arousal as all other non-traumatic stimuli (i.e., *ϕ*_*emot*_ = 1.0). Since during a treatment session, patients’ negative affect tends to increase in anticipation of the imaginal exposure exercise, we introduced the positive stimulus at a position spaced three stimuli prior to re-encoding the trauma memory. That way, positive affect decreased as the moment of re-encoding the trauma memory approached.

We then introduced two additional control conditions to clarify the mechanisms of potential change in the Imaginal Exposure condition. In the Positive Events control condition, we introduced a new, positive stimulus every 20 lists during the Neutral Period, at exactly the same locations in time at which virtual subjects experienced the positive stimuli prior to treatment in the Imaginal Exposure condition (described above). This condition verified that the Imaginal Exposure condition did not reduce negative emotion simply due to the positive emotional context that preceded each re-encoding of the traumatic memory. The properties of each positive stimulus matched those of the positive event in the Imaginal Exposure condition.

Finally, during a *Re-encoding only* control condition, virtual subjects reencoded the traumatic memory as a new encoding event, at the same five regular intervals during the Neutral Period, again spaced 20 lists apart. Unlike in the Imaginal Exposure condition, there were no positive stimuli to evoke a less-negative context prior to these re-encoding sessions. This modeled an ongoing question regarding the mechanisms of imaginal exposure treatment: specifically, if approaching experiences of the traumatic memory will lead to therapeutic change, why do patients not recover by reexperiencing the trauma-memory during flashbacks? Specifically, this condition tested whether simply re-encoding the memory in new contexts reduces negative emotional responding to the memory and promotes mood recovery, regardless of the emotional content of those new contexts.

#### Results

The four conditions introduced during the neutral period allowed us to assess whether CMR3 predicts reduced negative emotional responding to trauma memories: (1) spontaneously, with the passage of time and neutral events (Baseline Condition); (2) in response to new positive events (Positive Control Condition); (3) due to re-activating and re-encoding the memory, but without any changes in the patient’s current emotional context (Re-encoding Condition); or (4) upon re-encoding the trauma memory in a safe emotional context, as might occur in a therapy session (Imaginal Exposure Condition).

Figure 12 demonstrates the CMR3-predicted mood for an individual virtual subject, and Figure 13 shows CMR3-predicted mood aggregated across virtual subjects. During the Baseline Condition, trauma-memory intrusions continued to evoke high levels of negative emotion throughout the Neutral Period (Figure 12a; Figure 13). During the Positive Control Condition, positive stimuli briefly instated a positive emotional context in the moment. However, trauma-memory intrusions continued to evoke negative emotional context, thus returning mood to a negative baseline each time (Figure 12b; Figure 13). When virtual subjects reactivated the trauma memory and then reencoded it in association with less-negative contexts, each subsequent reactivation of the trauma memory evoked progressively reduced negative emotion, until mood stabilized near a neutral value of 0.0 (Figure 12c; Figure 13). Simply re-encoding the trauma memory, without altering the contents of its associated emotional context, did not result in mood improvements (Figure 12d, Figure 13).

**Figure 12.**
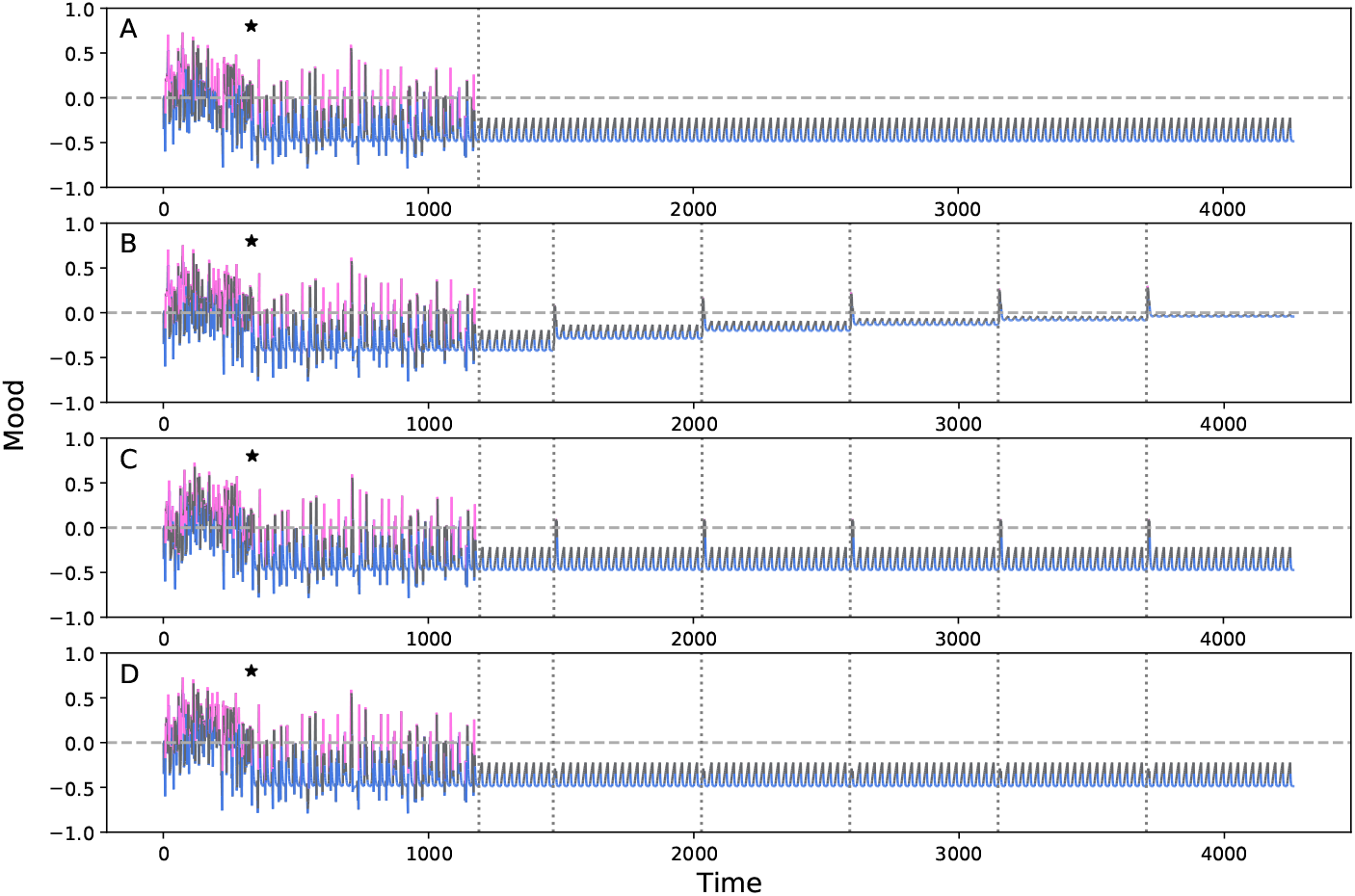
Imaginal Exposure Therapy Alleviates Model-Predicted Intrusive Memories. Across two simulation periods, delineated by a vertical line, one virtual subject experienced and remembered positive (pink), neutral (gray), and negative (blue) events. The vertical axis indicates model-simulated mood, ranging from -1 (max. negative) to -1 (max. positive). During an early Developmental Period, this virtual subject experienced 30% negative, 30% positive, and 40% neutral events, with one high-arousal negative event (traumatic event) at the location of the asterisk (*). Next, this virtual subject experienced a Neutral Period of unique, neutral events. This subject then experienced the following intervention conditions, in each of four distinct simulation runs. **A**. No intervention during the Neutral Period (Baseline Condition). **B**. Five sessions of reencoding the trauma memory in less-negative contexts, marked by vertical dotted lines (Imaginal Exposure Condition). **C**. Five encounters with positive stimuli, at the same timepoints as the imaginal exposures (Positive Control Condition). **D**. Five sessions of reencoding the traumatic event, with no change in emotional context prior to reencoding (Reencoding-Only Control Condition). *T*_1_-*T*_5_ = each of the five timepoints at which this virtual subject experienced the given intervention.

**Figure 13.**
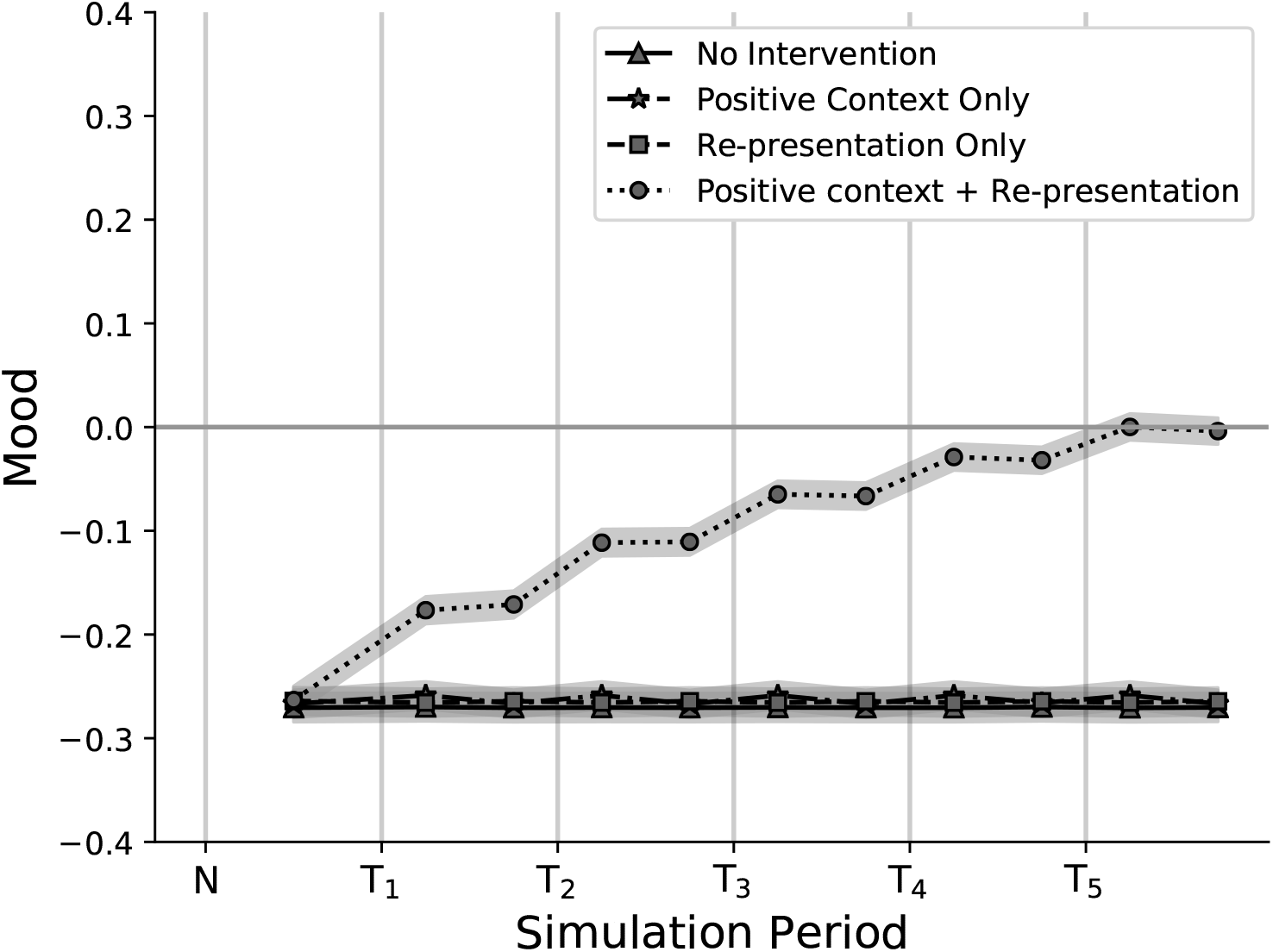
Effects of Trauma Exposure and Imaginal Exposure Therapy on Mood. Model-predicted mood for each of four conditions, aggregated across virtual subjects. Virtual subjects experienced a Developmental Period, containing an even mixture of negative and positive life events (30% negative, 30% positive), with one high-arousal negative event (*ϕ*_*emot*_ = 50.0). Next, virtual subjects experienced a Neutral Period of unique, neutral events, containing one of four distinct intervention conditions, in each of four simulation runs. Triangle markers indicate no intervention during the Neutral Period (Baseline Condition). Circle markers indicate five sessions of imaginal exposure therapy (bounded by vertical dotted lines), in which virtual subjects re-encoded the traumatic memory in a less-negative emotional context evoked by prior positive stimuli (Imaginal Exposure Condition). Star markers indicate five encounters with positive stimuli at the same times as the imaginal exposures, every 20 lists (Positive Control Condition). Square markers indicate five instances of reencoding the high-arousal negative event, but without any change in emotional context prior to reencoding (Reencoding-Only Control Condition). Vertical lines mark the beginning and end of each intervention session. The points between these lines represent the predicted mood during the first and second half of the session, respectively (averaging across each timepoint individually is not possible due to the stochastic nature of the simulation). N = The Neutral Period after the traumatic event has occurred. *T*_1_-*T*_5_ indicate the five timepoints at which a virtual subject receives the given intervention for that condition.

The rate of trauma-memory intrusions remained essentially constant over time, with minimal variability across virtual subjects. Therefore, we report the mean frequency of trauma-memory intrusions across the timepoints in each condition: the mean frequency was 13.8 (SEM = .003) for the Baseline and Positive-Events simulations and 14.0 (SEM = .020) for the Re-Encoding and Imaginal Exposure simulations. No condition decreased the frequency of trauma-memory intrusions, suggesting that imaginal exposure in PE may primarily serve to reduce negative affect associated with the trauma-memory, whereas *in vivo* exposure to trauma-cues may serve to reduce the frequency of trauma-memories’ intrusions, as observed in Simulation 6.

In the Imaginal Exposure Condition, the classic finding of reduced negative emotion, or apparent habituation, across rather than within sessions of imaginal exposure (Jaycox et al., 1998; M. Craske et al., 2008) emerged naturally from the model (Figure 12b; Figure 13). In CMR3, the trauma memory reinstates its old associated context when first retrieved, thus evoking strong negative emotion. Within the session, the memory forms associations to the new safe context, updating the trauma network in the patient’s episodic memory. However, the memory will not evoke its updated context until the next time the memory is retrieved, resulting in reduced negative affect across, but not within, sessions. By proposing that this phenomenon occurs due to updating in the episodic memory system, CMR3 predicts that imaginal exposure in PE can also reduce negative trauma-associated emotion across sessions for patients whose primary trauma-associated emotion is an emotion other than fear, such as guilt, shame, or grief (Brown et al., 2019).

## General Discussion

We propose a retrieved-context theory of how mnemonic processes contribute to mood and emotional disorders. At the core of this theory is a dynamic representation of context that organizes memories of life’s events based on both their time and place of occurrence, and on the perceptual, semantic and emotional attributes surrounding the event. Context associates with the features of new memories as they form, such that remembering an event will tend to reactivate its previously associated contexts. These retrieved contextual states drive the evolution of context, incorporating all past associations of a given experience into the current context, which in turn associates with subsequent experiences. These recursive contextual dynamics allow memories of temporally disjoint events to become linked through time owing to their shared contextual features (see, Kahana, 2020 for a review).

Following Talmi et al (2019), our implemention of retrieved-context theory (termed CMR3) assumes that context includes the emotional attributes of an experience. However, rather than representing emotion as synonymous with arousal (Talmi et al., 2019), we model distinct mood states in emotional disorders by differentiating positive and negative emotions. We further extend the model by enabling the accrual of prior experiences across a person’s full lifespan (Lohnas et al., 2015). This enables our CMR3 model to capture the dynamics of unwanted, intrusive memories from long-ago life contexts, as in PTSD.

Consistent with efforts to ground clinical theory in basic cognitive processes (Insel et al., 2010; Sanislow et al., 2010), our model does not predict the development of a *category* of disorder, such as Major Depressive Disorder (MDD) or Posttraumatic Stress Disorder (PTSD). Rather, our model describes the underlying basic processes in episodic memory that generate negative mood and memory disturbances, as transdiagnostic symptoms.

### Mood-congruent and Emotion-state Dependent Recall

We first show how CMR3 predicts the emotional clustering of episodic memories. Subjects tend to recall emotional items in same-valence clusters, such that recalling a negative item leads to subsequently recalling another negative item, and recalling a positive item leads to subsequently recalling another positive item (see Experiment 1, and also Siddiqui & Unsworth, 2011; Long et al., 2015). In CMR3, clusters of same-valence memories arise during recall because, for example, recalling a negative item evokes negative emotional context, which then cues the recall of other items associated to negative contexts. This arises because the similarity between encoding and retrieval contexts guides recall. Owing to this same principle, CMR3 also predicts both heightened recall of memories whose emotional properties match the current mood state (Mood-congruent Recall; Simulation 3), as well as heightened recall of even neutral memory material if it was encoded against an emotional backdrop that matches the person’s current emotional state (Emotion-State Dependent Recall; Simulation 4). That is, neutral memory content will be better recalled if the emotional context present just prior to and during encoding matches the emotional context present during retrieval.

### Persistent Negative Mood

Retrieved-context theory predicts that emotional context can linger after an event and bind to new events that follow, even if these new events are neutral (Simulation 4). Thus, negative emotional context can generalize throughout the memory network and reactivate when recalling neutral, or even positive, events. Increasing the ratio of positive to negative events, as in Behavioral Activation Therapy (Beck et al., 1979; Jacobson et al., 1996; Lewinsohn et al., 1980), evokes positive contexts that can also bind and generalize throughout the memory system, thus improving patients’ mood (Simulation 4, Treatment Period).

However, for patients experiencing intrusive negative memories, CMR3 predicted that positive event scheduling will briefly evoke positive emotion, but that the memory-intrusions will persistently reactivate their associated negative contexts, eventually blunting responsiveness to positive events (Simulation 5). Thus, our model corroborates the importance of targeting intrusive memories in treatments not only in PTSD, but also alleviating for negative mood in major depression (C. R. Brewin et al., 2009).

Encoding an event in multiple contexts associates its memory with a larger number of potential retrieval cues (Lohnas & Kahana, 2014). Thus, repeatedly experiencing the same stressful event, or ruminating over the event in varied contexts, will increase the retrieval routes to that memory and thus its likelihood of spontaneous subsequent recall (Simulation 5). Indeed, rumination, which includes the repetitive rehearsal and elaboration upon negative memories, is associated with and can prompt intrusive memories in PTSD (Michael et al., 2007).

### Intrusive Memories

Intrusive memories of negative events occur prominently in PTSD, but also across other emotional disorders (C. Brewin, 2010), and for painful events not traditionally categorized as traumatic (Brozovich & Heimberg, 2008; A. D. Williams & Moulds, 2007). Retrieved-context theory proposes a general mechanism for intrusive memories, in which encountering stimuli whose features resemble the perceptual, spatial, and temporal context that directly preceded a negative event will cue intrusive memories of that event (Simulation 6), even if the stimuli have no semantic relationship to the event, as is common in PTSD (Ehlers & Clark, 2000; Ehlers et al., 2004). Similar to Talmi et al. (2019), we propose that high emotional arousal during a traumatic event leads to stronger encoding of associations between the traumatic memory and its context. These strengthened contextual associations increase the probability that unrelated contexts will trigger the memory.

Trauma-memory intrusions often occur with vivid imagery and a sense of “nowness,” i.e., that the memory is happening again in the here-and-now (Hackmann, Ehlers, Speckens, & Clark, 2004). Retrieved-context theory conceptualizes a memory’s vividness as how strongly the retrieval context activates that memory’s features (Simulation 7). Drawing on Tulving’s conception of mental time travel (Tulving, 1983), we model the “nowness” of a memory as the similarity between the memory’s encoding context and the context present at the time of retrieval (Simulation 8). However, mental time travel is but one possible framework for perceived “nowness.” A different interpretation is that nowness involves a fading of the current context into the background, not a sense that the intrusion belongs in the current context, as we propose.

Although our simulations focused on intrusive negative memories, CMR3 can also predict spontaneous intrusions of high-arousal positive memories, consistent with empirical findings (Berntsen, 1996; Berntsen & Rubin, 2002). Simulations 6-9 set *ϕ*_*emot*_ to modulate context-to-memory associations only for negative events, in order to examine the effects of high-arousal negative events without “background noise” from the effects of high-arousal positive events. However, future work setting *ϕ*_*emot*_ to also modulate contextual associations for positive events would result in enhanced memory for, and spontaneous intrusions of, high-arousal positive memories as well.

### Treating Intrusive Trauma Memories

Our model suggests two treatment routes for trauma-memory intrusions. The first route is modifying the context-to-memory associations in the emotion network, as in Prolonged Exposure Therapy (PE; Foa & Kozak, 1986; Foa et al., 2007). Associating trauma cues to neutral or non-traumatic contexts, as in *in vivo* exposure, increases the probability that these stimuli will cue nontraumatic memory content, and reduces the frequency of negative memory-intrusions (Simulation 6). Further, reactivating and then reencoding the trauma-memory in association with new, safe or neutral contexts as in *imaginal exposure*, even if the trauma-memory itself evokes strong negative affect against the backdrop of these less-negative contexts, decreases negative affect across sessions (Simulation 9).

The second route for alleviating intrusive memories involves modifying a patient’s current contextual state to reduce its overlap with the trauma-memory’s encoding context. Reduced negative emotion due to cognitive restructuring would reduce the tendency of mood-congruent recall processes to reactivate the trauma-memory and other negative memories (Simulation 2). This might occur through restructuring maladaptive cognitions and thereby reducing shame and guilt, as in Cognitive Processing Therapy (CPT; Resick & Schnicke, 1992), as well as during processing in PE.

### Individual Differences in Risk and Resilience

Retrieved-context theory proposes specific episodic memory processes that serve as risk or protective factors. Fitting our model to a subject’s memory patterns can identify the parameters that determine how strongly a given memory process is in play for an individual subject.

#### Emotional Valence

The *β*_*emot*_ parameter governs how quickly a subject’s internal context updates to take in new emotional content. Swift emotional updating increases the level of emotional context that is present in a subject’s internal contextual state, and thus, the level of emotional context present to associate with new memories as they form. When negative events occur, high values of *β*_*emot*_ may increase the levels of negative emotion present in subjects’ daily context, resulting in more negative mood states (Simulation 7).

#### Emotional Arousal

Negative arousal strengthens context-to-memory associations during the encoding of negative memories. Thus, higher values of *ϕ*_*emot*_, the scalar that modulates these associations, will enhance recall of this material and predispose trauma- and other negative memories to intrude spontaneously into unrelated contexts (Simulations 6-9).

#### General Parameters

Model parameters that address general memory function can also help to explain individual differences that predispose a person to develop psychopathology. For example, *c*_*thresh*_ controls the degree to which subjects can exert cognitive control to filter and censor unwanted memory content, such as trauma-memory intrusions.

### Relation to Existing Theories

#### Recalling Trauma in PTSD: One Memory System or Two?

Our model speaks to the longstanding controversy over whether more than one memory system is needed to account for the way in which traumatic events are recalled (C. R. Brewin, 2014; Bisby, Horner, Bush, & Burgess, 2018; Bisby, Burgess, & Brewin, 2020; Rubin et al., 2008, 2011).

Retrieved-context theory proposes that trauma memories and other autobiographical memories arise from common factors, within a unitary memory system. As in the Autobiographical Memory Theory of PTSD (Rubin et al., 2011), our model proposes that emotional intensity, repetition, and rehearsal strengthen the accessibility of a memory and thereby predispose it to spontaneously activate. In CMR3, contextual associations in episodic memory serve as the core process underlying these principles – heightened emotional intensity strengthens context-to-item associations (Simulations 6-9, and see Talmi et al., 2019); repetition and rehearsal associates the memory to a wider array of contexts that can serve as retrieval cues (Simulation 5, and see Lohnas, Polyn, & Kahana, 2011; Madigan, 1969; Melton, 1970); and the *semantic contiguity effect* generates heightened recall of memories that have strong semantic relations with other memories and with the current context (Simulation 1, and see Howard & Kahana, 2002b), potentially enhancing recall for memories that have meaningful relatedness to the person’s life story.

Dual-representation theory (DRT) proposes that two memory systems are necessary to explain memory recall in PTSD. Both systems are thought to be part of normal healthy memory but to behave differently under extreme stress. Specifically, DRT proposes that high-arousal negative emotion at encoding modulates the strength of associations between traumatic memories and their encoding contexts (C. R. Brewin, 2014; C. R. Brewin et al., 2010). Thus, high arousal negative emotion disrupts the associations between events’ perceptual features and their spatiotemporal contexts (C. R. Brewin, 2014; Bisby et al., 2018). As a result, DRT proposes that trauma memories have unique properties that differ from those of other nontraumatic negative episodic memories. Further, DRT proposes that reunification of the memory trace – reassociating events’ perceptual features to their contextual elements – is needed to reduce intrusive memories in PTSD (C. R. Brewin, 2014).

When high-arousal negative emotion occurs, particularly in the absence of dissociation, which both theories predict would interfere with new learning, dual-representation and retrieved-context theories predict different outcomes for exposure-based treatments. During exposure exercises, patients face feared stimuli, including feared memories, and are encouraged to fully experience their distress. Heightened levels of negative emotion and negative physiological arousal during exposures are either associated with better treatment outcomes (M. Craske et al., 2008; Kircanski et al., 2012; A. J. Lang & Craske, 2000; Jaycox et al., 1998), or else have no relation with treatment outcomes (Rupp, Doebler, Ehring, & Vossbeck-Elsebusch, 2017). In retrieved-context theory, high-arousal negative emotion during exposure is beneficial, by more strongly binding the memory to the new, therapeutic context.

In dual-representation theory, high-arousal negative emotion disrupts the binding of items-to-contexts during new learning, even for non-personally relevant stimuli in a laboratory setting (Bisby et al., 2018). Accordingly, the negative emotion present during an even more emotionally evocative, imaginal exposure to the memory of prior trauma would be an unhelpful, or even harmful, byproduct of the treatment’s true active ingredient: re-unifying the memory trace. Dual-representation theory predicts that Prolonged Exposure Therapy for PTSD and other exposure-based therapies will still be efficacious, as long as the high-arousal negative emotion is carefully modulated during session, thus preventing it from interfering with the goal of re-unifying the reactivated trauma memory (Bisby et al., 2018).

Supporting dual-process theory, laboratory studies have found weaker associations during recall between negative and neutral stimuli than between pairs of neutral stimuli (Bisby & Burgess, 2014; Bisby et al., 2018). However, in Experiment 1, we replicated the reduced tendency of emotional items to cue subsequent recall of differently-valent items as part of the *emotional clustering effect*. We demonstrate that our model predicts this effect through the mechanism of emotional context similarity (Simulation 1).

In addition, Bisby and Burgess (2014) and Bisby et al. (2018) did not control for greater semantic relatedness among pairs of negative (vs. pairs of neutral) stimuli (Talmi & Moscovitch, 2004). Consequently, this pattern of results may be accounted for by the classic “fan effect” (Anderson, 1974). Because neutral items have weaker semantic associations to other neutral items, as well as to items of negative or positive valences, the associations that neutral items form to other neutral items during the experiment will have less competition from prior, long-standing associations. As a result, negative items have stronger pre-experimental associations to other negative items that will compete for retrieval, and thus impair recall, of new associates learned during the experimental context. Nonetheless, this explanation is just one of many possibilities, and further empirical testing is needed to actually establish greater semantic relatedness of negative items as the underlying mechanism in Bisby and colleagues’ work.

#### Emotional Processing Theory

Emotional processing theory (EPT) has generated a highly efficacious treatment for intrusive trauma memories in PTSD called Prolonged Exposure therapy (PE; Foa & Kozak, 1986; Foa et al., 2007; Powers et al., 2010). In EPT, the trauma-memory exists in a network of associations with other stimuli, maladaptive cognitions, and fear responses, called the *fear network* or *emotion network* (Foa & Kozak, 1986; Brown et al., 2019; Lane, Ryan, Nadel, & Greenberg, 2015). Treatment reactivates the memory in order to restructure this network, by exposing the patient to the feared stimulus (in imaginal exposure, this is the memory itself). Retrieved-context theory shares several core principles with EPT. In CMR3, fully reactivating the trauma memory is necessary to restructure its network of contextual associations, thereby reducing the frequency of trauma-memory intrusions (Simulation 6) and the patient’s distress upon recalling the trauma-memory (Simulation 9).

Retrieved-context theory differs from EPT regarding the role of habituation. EPT arose from early models of fear conditioning, including the classic Rescorla-Wagner model (Rescorla & Wagner, 1972). In the Rescorla-Wagner model, presenting the feared stimulus in the absence of the feared outcome weakens and ideally breaks the stimulus-response relationship (Rescorla & Wagner, 1972). Accordingly, EPT proposes that staying present with the stimulus until habituation occurs within the session weakens the association between the stimulus and the conditioned fear response, such that the memory no longer activates the fear response.

In PE, within-session habituation to the trauma memory does not reliably reduce symptoms (Rupp et al., 2017; Jaycox et al., 1998). Rather, across session habituation appears more strongly associated with symptom reduction. That is, PE patients often experience extreme distress while reactivating the trauma-memory, find that their distress does not reduce within the session, and yet later experience reduced distress when reactivating the trauma memory in a new session. This finding has raised concerns about circularity: perhaps “those who habituate, habituate,” or habituation is incidental to the success of exposure-based treatments (C. R. Brewin, 2006).

To address these issues, recent updates to EPT incorporate principles of inhibitory learning, proposing that exposure-based treatments introduce new, competing CS-noUS associations, rather than breaking the preexisting CS-US association (Brown et al., 2019). Inhibitory learning and the effects of expectation violations enhance and powerfully contribute to new therapeutic learning when present (M. G. Craske, Treanor, Conway, Zbozinek, & Vervliet, 2014; M. G. Craske, Liao, Brown, & Vervliet, 2012; Brown, LeBeau, Chat, & Craske, 2017).

Retrieved-context theory offers an alternative mechanism of restructuring the episodic memory network. In CMR3, recalling the memory first reactivates its old emotional context, and then re-encodes the memory in association with a new, less-negative emotional context. The newly updated emotional context is only reactivated the next time that the patient retrieves the memory, such that negative affect associated with the trauma memory decreases across, rather than within, sessions. As a result, reduced negative emotion across, rather than within, exposure sessions emerges naturally from our model (Simulation 9).

Within an individual session of PE, the trauma memory may be rehearsed multiple times. In CMR3, repeating the trauma-memory within sessions will have a weaker effect on the trauma-memory network than repetitions across sessions, because context remains relatively constant within an individual session. This is a clinical application of the *spacing effect*, in which spaced rather than massed repetitions enhance learning by associating repeated experiences with more diverse contexts (Lohnas & Kahana, 2014). A further consideration is that, although EPT is grounded in fear-conditioning principles, PE is still highly efficacious in treating PTSD for patients whose primary emotional response to their traumatic event involved an emotion other than fear, such as shame, guilt, or sadness (Brown et al., 2019), which is common for PTSD patients (Reynolds & Brewin, 1999; Resick & Schnicke, 1992). Our model proposes that in episodic memory, the emotional context associated with an episodic memory trace can incorporate an amalgam of different emotional responses, including but not limited to fear. Thus, reactivating and reencoding the trauma’s episodic memory trace in association with novel contexts would still generate the clinical pattern of heightened negative emotion within a session, but reduced negative emotion in subsequent sessions, even for non-fear emotions. Accordingly, our model proposes a generalized framework, in which heightened negative arousal during encoding strengthens a memory’s contextual associations, and thereby results in intrusive memories of negative events in patients with depression and other emotional disorders, not just PTSD (C. Brewin, 2010).

Overall, retrieved-context theory generates several novel predictions for exposure-based treatments. First, even when patients confirm their expectations that, upon reactivating a trauma-memory, they will both experience extreme distress and even tolerate that distress poorly, repeated exposures in positive or less-negative emotional contexts will still result in symptom improvement. Second, the duration of within-session exposure to the memory is not the primary driver of reduced negative affect, as long as the memory is fully reactivated for re-encoding, and the background context for reencoding is sufficiently “non-negative” emotionally (note that emotional content of the surrounding reencoding context, such as the emotional tone of the therapist’s office, is distinct from the emotion evoked by the trauma-memory). Third, PE therapy will still alleviate trauma-memory intrusions even when the primary emotion associated with the trauma is not fear. Finally, the PE principles of reactivating and reencoding distressing memories, originally pioneered for the treatment of trauma memories in PTSD (Foa & Kozak, 1986), may also alleviate distress and memory-intrusions resulting from painful events in depression and other disorders.

#### Spreading Activation Theory

Bower (1981) proposed that mood-congruent and emotion-state dependent memory occur because emotional valence serves as a node in semantic networks. A positive or negative emotion thus activates semantically associated memories, which tend to share the same emotional properties due to semantic relatedness. However, in retrieved-context theory, because emotional valence also serves as a part of a memory’s episodic context, it can become associated with stimuli that themselves are neutral and have no semantic relation with negative emotion. This allows our theory to model the generalization of negative affect to neutral events (Simulation 4). Emotional context also allows our model to account for how unrelated stimuli immediately preceding a traumatic event can later cue intrusive memories and trauma-associated affect (Simulation 6, and see Ehlers & Clark, 2000).

#### Reconsolidation

Reconsolidation theory suggests that imaginal exposure improves PTSD symptoms by updating and re-encoding the trauma memory to include new emotional elements (Lane et al., 2015). Our account of imaginal exposure does not involve a memory reconsolidation process. Rather, in CMR3, reactivating and reencoding traumatic memories during imaginal exposures associates those memories with novel contexts. Accordingly, imaginal exposure can reduce negative affect without changing the contents of the traumatic memory itself.

## Limitations

We have intentionally pursued a highly simplified approach to investigating a problem of great complexity, namely the role of memory and emotion in the development and treatment of emotional disorders. Although learning and recalling lists of words clearly lacks the complexity of everyday experiences, this paradigm has allowed researchers to identify and quantify the core principles of human memory, including primacy, recency, temporal and semantic contiguity effects (Kahana, 1996; Murdock, 1967; Howard & Kahana, 2002b; Sederberg, Miller, Howard, & Kahana, 2010). These principles, in turn, underlie memory for more complex real-life events (Jansari & Parkin, 1996; Loftus & Fathi, 1985; Moreton & Ward, 2010; Uitvlugt & Healey, 2019; Healey et al., 2019). We view our model’s setup as a springboard for more realistic models of human experience.

Further research is needed to determine the precise relationship between arousal and the learning rate for new associations during encoding. One could easily introduce a non-linear relation between arousal and learning rate, as in the classic Yerkes-Dodson (1908) results, and perhaps this would be a fruitful direction for future work. Or, perhaps under extreme levels of arousal, there is a more systemic shutdown of the cognitive system, as suggested by dual-representation theorists.

Another limitation lies in our decision to model emotion as varying along dimensions of valence and arousal. This allows the model to account for patterns of comorbidity between anxiety, depression, and other emotional disorders, as well as for how intrusive memories can occur for many types of distressing events. However, emotion is a complex, multivariate construct. For example, anger and fear are both negatively valent and high-arousal emotions, and thus would be indistinguishable in the current CMR3 model, yet they have distinct subjective experiences, action tendencies (avoid vs. approach; Carver & Harmon-Jones, 2009), and neural patterns (Kragel & LaBar, 2015).In addition, not all negative emotions induce high arousal. For example, disgust may be associated with reduced physiological arousal (Ekman, Levenson, & Friesen, 1983). Computational approaches, such as the one we have undertaken with CMR3, will benefit from future empirical work elucidating the role of categorical features of emotion on memory, beyond the dimensional features of valence and arousal.

Further, processes of cognitive control play an important role in autobiographical memories. One of the key elements of CMR3 is its ability to use context-similarity as a filter to suppress memories that are contextually distant from the target of conscious awareness or attention, operationalized via the *c*_*thresh*_ parameter. This cognitive control mechanism serves a critical role in modeling the reactivation and filtering of trauma-memory intrusions. However, CMR3 does not fully specify all possible sources of cognitive control that subjects may use to guide their thought process. The companion question, to what extent are people able to control this memory process, remains an important area for future research.

Finally, a core phenomenon shared by both major depression and PTSD is overgeneral memory, or difficulty recalling specific memory episodes, likely related to or perhaps a consequence of deficits in context discrimination abilities (J. M. G. Williams et al., 2007). Our simulations do not directly address this phenomenon, but the CMR3 model offers a novel potential framework. In CMR3, experiencing repeated negative events, or multiple negative events that all share some common categorical information, across varied temporal contexts, will tend to form a web of associations across lists, much as in an experiment in which subjects experience repeated items from the same categories across lists (e.g., Postman, 1976; Anderson & Bower, 1972). Such manipulations cause high levels of associative interference. If the negative categories constitute the repeated items, reactivation of these categories will lead to higher levels of recall, whereas associative interference will lead to lower levels of event-specific recall. This is analogous to the effects of repetition in word lists (Zaromb et al., 2006; Miller, Weidemann, & Kahana, 2012). Although this mechanism could produce overgeneral memory in depressed patients, a proper modeling exercise would require experimental data in which depressed patients study lists of category-exemplar pairs of varying valence/arousal and show improved memory for the categories but impaired memory for the examples. We suggest this as an interesting direction for future work.

## Conclusion

Retrieved-context theory proposes a new computational, transdiagnostic model of memory and mood in emotional disorders. Together with the emerging field of computational psychiatry (Bennett, Silverstein, & Niv, 2019; Montague, Dolan, Friston, & Dayan, 2012), we propose that clinical theory will benefit from testing whether, for an appropriate set of inputs, a formal model generated from a given theory can predict the behavioral patterns observed in laboratory and clinical settings. The resulting model should be able to predict the development of symptoms, their maintaining factors, and the efficacy of existing treatments. Additionally, the model’s mechanisms should map on to literature regarding human cognition and neural architecture. Our simulations link clinical findings with cognitive operations, laying the groundwork for future work connecting these processes with neural mechanisms. We hope this framework will be used to both refine and challenge our model, thus furthering understanding of memory and mood.

## Appendix

Parameter Estimation and Model Comparison Methods We used a particle swarm optimization method to fit each model (CMR2, eCMR and CMR3) to each individuals’ behavioral data. Like a genetic algorithm or downhill simplex technique, a particle swarm searches a high-dimensional parameter space to find the optimal value of some objective function – in our case, the objective function is minimizing the error between predicted and observed data. Each particle in a swarm refers to a set of parameters (we used 200 particles) for a given model. We evolved the particles over 30 iterations, searching for the parameter set that provided the best fit to the behavioral data, separately for each model (CMR2, eCMR, and CMR3) and for each subject. In a given “run” of the model with one “particle,” we simulate an entire 24-session experiment, using the actual list items that a given subject experienced to determine the sequence of items presented to the model. The model then generates a set of simulated recall sequences, which we analyze to estimate the same behavioral patterns shown in Figure 2.

Having generated simulated behavioral data obtained for a given particle for one of our three models, we can now calculate a goodness-of-fit statistic quantifying the deviation between model and data. For our fit index, we used the *χ*^2^ error, defined as the sum of squared residuals, with each squared residual normed by the unbiased sample variance of the corresponding data point (Bevington & Robinson, 2003). After the *χ*^2^ error for each particle is determined, the optimal parameter set is identified as the one having the lowest *χ*^2^ error in this iteration. The particle-swarm algoirthm then adjusts the other particles to produce results similar to the current optimal value (hence the term “swarm”). Over the course of iterations, the particles converge on an optimal set of parameters, thereby minimizing the *χ*^2^ error.

The 56 points contributing to the *χ*^2^ error for each model fit comprised the first 15 and last three positions of the serial position curve (Fig 2a); the first 15 positions of the probability of first recall curve (Fig 2b); the lag-CRP curve at lags of -3 through +3; the nine conditional probabilities of transitioning among negative, positive, and neutral item types (negative-to-negative, negative-to-positive, negative-to-neutral, and so forth); all six points on the semantic CRP curve; the frequency of PLI’s per list; and the frequency of ELI’s per list.

Particle swarm optimization yielded best-fitting parameters for each of the 97 subjects in Experiment 1. At the end of the particle swarm process, the mean *χ*^2^ error across model fits to individual subjects was 36.60 (SEM = 4.95) for CMR2, 30.45 (SEM = 2.05) for eCMR, and 29.74 (SEM = 1.64) for CMR3, where lower *χ*^2^ error values indicate better model fits. Degrees of freedom for the *χ*^2^ error values were determined by n – m, where n equals the number of data points fit (56 points) and m equals the number of free parameters in each model (15 for CMR2 and 17 for eCMR and CMR3). Thus, for 97 subjects, CMR2 provided 86 model fits with non-significant *χ*^2^(41) error; eCMR provided 92 model fits with non-significant *χ*^2^(39) error; and CMR3 provided 90 model fits with non-significant *χ*^2^(39) error. Table 2 presents the means and standard errors of the sets of parameters obtained for CMR2, eCMR, and CMR3, taken across individual-subject fits.

After obtaining the best-fitting parameters for each model (Table 2), we used these parameters to simulate a full dataset of 24 sessions of delayed free recall for each model. To assess each model’s ability to fit the aggregate data, we calculated three measures of fit: (1) the same *χ*^2^ goodness of fit index that was minimized while fitting the model; (2) the Bayesian Information Criterion (BIC) to account for the different number of parameters across models (Kahana et al., 2007; S. M. Polyn et al., 2009; Schwarz, 1978); and (3) the RMSE, to identify which specific behavioral analyses determined each model’s overall ability to fit the data. We calculated each goodness-of-fit test with respect to all data points in all analyses, *n* = 75, to obtain a total measure of each model’s ability to fit the aggregate behavioral effects.

The resulting *χ*^2^ error values were *χ*^2^(60) = 74.1, *p* = .10 for CMR2, *χ*^2^(58) = 55.2, *p* = .58 for eCMR, and *χ*^2^(58) = 56.7, *p* = .52 for CMR3, indicating that all three model fits had non-significant error. For direct model comparisons, it is not valid to directly compare the size of *χ*^2^ error values because eCMR and CMR3 have two more parameters than CMR2, which gives them an advantage over CMR2 in their ability to fit the data. Therefore, we calculated the Bayesian Information Criterion (BIC) values (Kahana et al., 2007; S. M. Polyn et al., 2009; Schwarz, 1978) for each model’s fits to the aggregate data. The BIC accounts for each model’s ability to fit the data while penalizing models that have a greater number of parameters, thus placing the three models on equal footing for comparison. Under the assumption of normally distributed residuals, the BIC formula simplifies to:

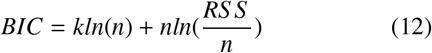

Here, *k* represents the number of free parameters in the model, *n* represents the number of data points, and RSS is the residual sum of squares. To ensure that all points contribute equally to model fits, we multiplied the emotional clustering effect by a factor of 10 to place it on the same scale as the other data points from the set of behavioral analyses (S. M. Polyn et al., 2009). This was not necessary for the *χ*^2^ error values because norming the squared residuals by the unbiased sample variance in the data (Bevington & Robinson, 2003) already sets the contributing residuals to comparable scales. The resulting BIC’s were -347.06 for CMR2, - 345.65 for eCMR, and -353.82 for CMR3, where lower (i.e., more-negative) values represent improved model fits. The results indicate that CMR3 provided the best balance of model parsimony and error minimization, followed by CMR2 and then eCMR.

To identify which behavioral effects distinguished each model’s ability to fit the aggregate data, we calculated RMSE values for each behavioral analysis (Table 3). For the total RMSE, taken across all points in all graphs (N = 75), CMR3 provided the smallest RMSE, followed by CMR2, and then eCMR, where smaller values indicate better model fit. Comparing eCMR and CMR3, eCMR provided lower RMSE’s for the positive lags of the Lag-CRP and the frequency of extra-list intrusions. Conversely, CMR3 provided the lowest RMSE for the emotional clustering effect, followed by CMR2 and then eCMR. CMR2 provided worse fits to the semantic clustering effect and the probability of first recall, suggesting that the model may have had to sacrifice fits to these data in its attempts to capture emotional clustering patterns.

